# Heterologous Rieske non-heme iron monooxygenases enable efficient microbial conversion of lignin guaiacol to adipic acid

**DOI:** 10.64898/2026.02.23.707458

**Authors:** John F.C. Steele, Charles B. Wackwitz, Gary Walker, Krishnan T. Selvy, Stephen Wallace

## Abstract

Adipic acid (1,6-hexanedioic acid) is a key building block for nylon-6,6, a widely used polymer in the global chemical industry. Current industrial production relies on petrochemical feedstocks and nitric acid oxidation of cyclohexane/cyclohexanol mixtures, releasing nitrous oxide, a potent greenhouse gas. Biotechnological routes offer sustainable alternatives but have been limited by low yields or reliance on multi-strain systems. Here we report a one-pot, single-strain microbial process for the efficient conversion of guaiacol – a lignin derived aromatic – into adipic acid. By integrating heterologous Rieske non-heme iron monooxygenases from *Cupriavidus necator* N-1 with systematic process optimisations in engineered *Escherichia coli*, we achieve near-quantitative conversion with 97% yield and titres of 1.5 g/L in aqueous, lab-scale reactions. This work demonstrates a novel and efficient strategy for lignin valorisation through engineered microbial synthesis, providing a new sustainable and scalable route to adipic acid.

## Introduction

Modern chemical manufacturing faces two central challenges: dependence on finite petrochemical feedstocks and the release of greenhouse gasses during production. With ambitious targets to reduce carbon emissions, there is an urgent need for sustainable processes that can meet demands for commodity chemicals. Addressing these challenges has driven cross-disciplinary research, particularly in engineering biology, where biologists, chemists, and engineers are developing biotechnological routes to essential chemicals in more environmentally benign ways.

Adipic acid (1,6-hexanedioic acid) is a high-profile target for greener manufacturing as the co-monomer of nylon-6,6, a polymer produced annually at multi-million-ton scale. Current industrial synthesis involves nitric acid oxidation of cyclohexanone/cyclohexanol mixtures, releasing nitrous oxide – a greenhouse gas with >200-fold the global warming potential of CO_2_[1]. Alternative strategies to adipic acid have included catalytic oxidation of benzene derivatives[2], semi-biological pathways[3], and fully biological routes[4,5] in *Escherichia coli*, yeasts[5], and *Pseudomonas putida*[6,7]. Engineered microbes are particularly attractive as they eliminate nitrous oxide emissions while offering the flexibility to convert post-consumer and industrial waste materials into valuable products under mild reaction conditions[8–10].

Among potential waste feedstocks, lignin is especially appealing. As the second most abundant biopolymer on Earth, lignin is a renewable, non-food resource rich in aromatic subunits. Depolymerization yields compounds such as guaiacol, syringol, and 4-hydroxyphenol, which all represent promising starting points for bioconversion. Following lignin depolymerisation, syringyl (S units), guaiacyl (G units) and *p*-hydroxyphenylpropane (H units) phenylpropanoid monomers can be integrated into central metabolism of engineered microorganisms and redirected towards diverse functional biomolecules including terpenoids, alcohols, fatty acids and polyketides.

In species such as *Pseudomonas putida* KT2440, *Acinetobacter baylyi* ADP1[11] and *Amycolatopsis* sp. ATCC 39116[12], lignin aromatics – including guaiacyl and *p*-hydroxyphenylpropane units – are metabolised using native pathways converging on protocatechuic acid (PCA). PCA then enters the β-ketoadipate pathway and ultimately the tricarboxylic acid (TCA) cycle[13]. Genetic manipulation of these pathways enables accumulation of valuable intermediates: for example, deletion of aldehyde reductase *PP_2426* leads to vanillin production in *P. putida*, while targeting muconate isomerases results in accumulation of the bioprivileged compound *cis,cis-muconic* acid during guaiacyl metabolism[12,14]. Extensive metabolic engineering in the genetically tractable lignin metabolizer *P*. *putida* has enhanced aromatic utilisation through heterologous demethylase expression[15], metabolic funnelling towards single products[16,17], and enzyme evolution to expand substrate scope to otherwise inaccessible syringyl monomers[18,19]. Similar engineering strategies have been applied in non-native hosts, with catabolism of lignin-derived phenolics established in *Saccharomyces cerevisiae*[20] and extensively developed in *E. coli*[21].

We and others have previously demonstrated a one-pot microbial route from guaiacol to adipic acid via catechol, *cis,cis*-muconic acid (ccMA), and 2-hexenedioic acid intermediates (Figure 1)[7]. This cascade relies on three heterologous enzymes – guaiacol demethylase[22], catechol dioxygenase, and muconate cycloisomerase/reductase[23]. The formaldehyde byproduct generated in vivo is detoxified by the native glutathione (GSH)-dependent frmRAB pathway, regulated by FrmR, which coverts formaldehyde to formate via *S*-formylglutathione before regenerating GSH[24]. While these studies established proof-of-concept for biological adipic acid synthesis, yields and titres were limited by inefficient guaiacol demethylation, creating a bottleneck at the pathway entry point.

**Figure 1.**
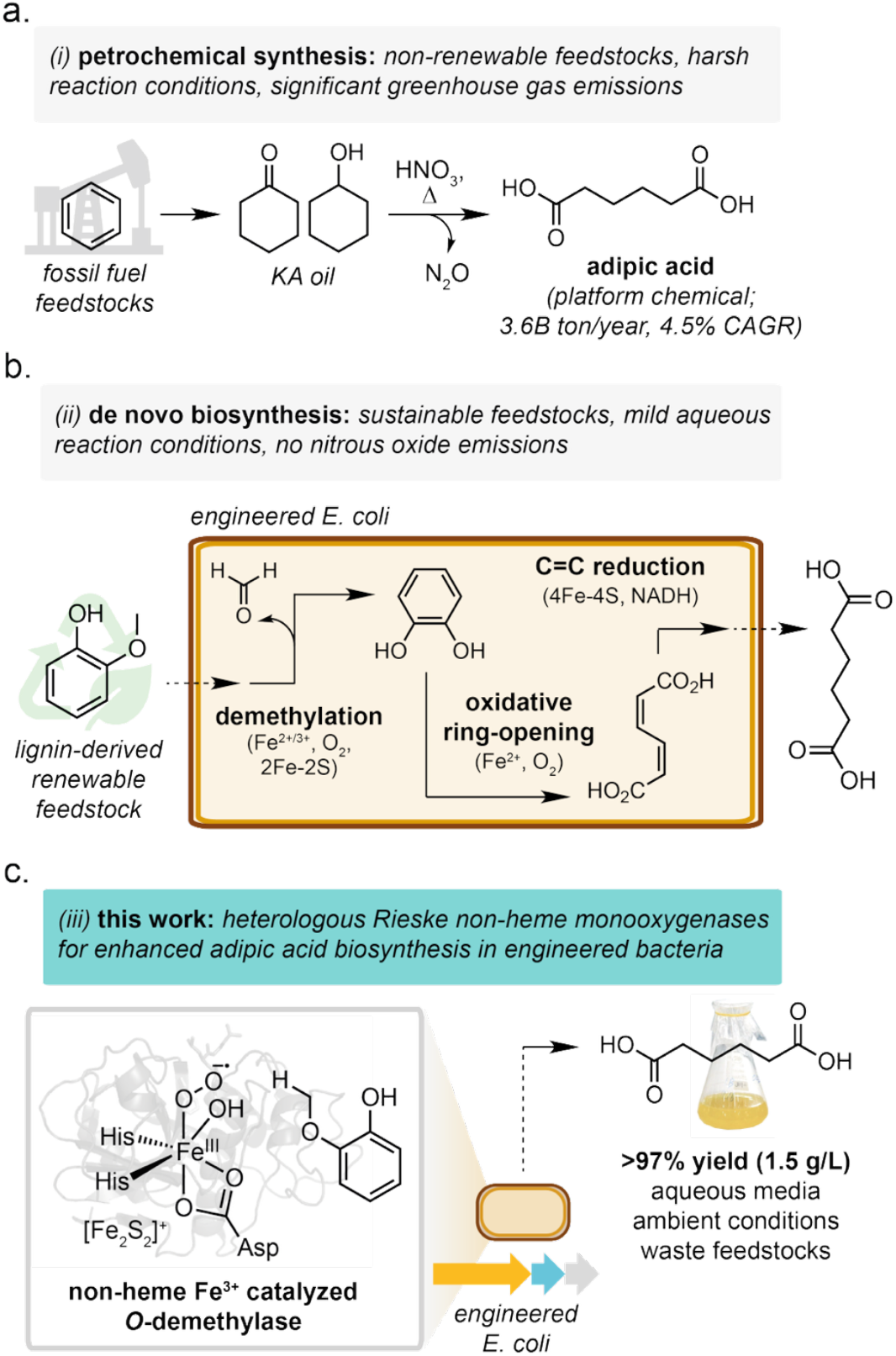
Overview of adipic acid production using current industrial and lignin valorisation routes. A) industrial production of adipic acid is performed by oxidation of petrochemical cyclohexanol/cyclohexanone mixtures with concentrated nitric acid, liberating stoichiometric nitrous oxide as a byproduct. B) Bioconversion of lignin-derived guaiacol to adipic acid proceeds by an initial demethylation, producing catechol and formate. Intradiol dioxygenation of catechol results in formation of ccMA, which can undergo sequential enzymatic reductions to produce adipic acid. C) The catalytic centre of non-heme rieske monooxygenases rely on a coordinated mononuclear iron to allow efficient guaiacol demethylation. Introduction of these alternative demethylases into engineered E. coli allows improved substrate conversion and increased adipic acid titres.

Here, we report a microbial system that overcomes this limitation by employing heterologous Rieske non-heme iron monooxygenases from *Clostridium* sp. Together with systematic process optimization in engineered *E. coli*, this strategy enables near-quantitative conversion of guaiacol from sustainable sources into adipic acid, achieving 97% yield and titres of 1.5 g/L under laboratory-scale conditions.

## Results and discussion

### Identification of efficient guaiacol demethylases

We previously demonstrated that *E. coli* BL21(DE3) can be engineered to convert guaiacol to adipic acid through a four-step pathway comprising guaiacol *O*-demethylation, catechol *ortho*-dioxygenation, and sequential reduction of *cis,cis*-muconic acid and 2-hexenedioic acid intermediates[8] (Figure 1). While this represented the first one-pot microbial route from guaiacol to adipic acid, titres were modest (3 mM, 0.44 g/L), and substantial substrate remained after whole-cell biotransformations.

To improve pathway efficiency, we targeted the initial *O*-demethylation step, reasoning that increased substrate turnover at the pathway entry-point would relieve a major bottleneck. Our previous system employed the guaiacol demethylase pair GcoAB from *Amycolatopsis* sp. ATCC 39116, consisting of a heme-dependent cytochrome P450 (GcoA) and its redox partner (GcoB), which transfers electrons from NADH via FAD and a 2Fe-2S cluster (Figure 2A) [25]. More recently, Rieske non-heme iron oxygenases have been identified that catalyze *O*-demethylation of guaiacol and related alkyl-catechols[26,27]. These enzymes use mononuclear iron rather than heme to activate molecular oxygen (Figure 2B, C). In both systems, substrate binding displaces water from the active site, enabling formation of a reactive Fe(IV)-oxo intermediate that drives demethylation through single-electron transfer and radical rebound chemistry.

**Figure 2.**
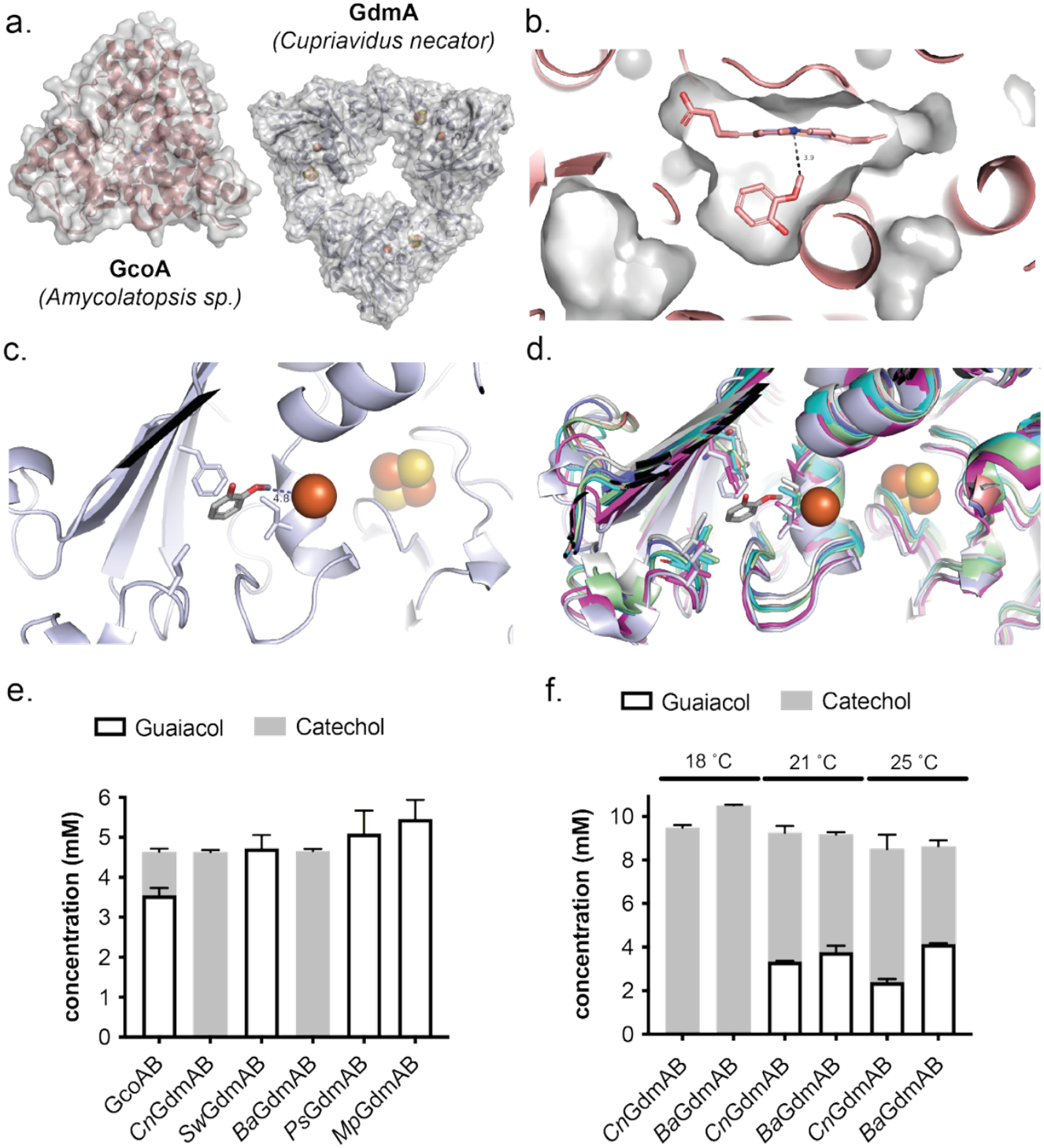
Screening candidate alternative demethylases A. Global fold of heme-P450 GcoA used in benchmark system, PDB 5NCB, and CnGdmA as a trimer, generated using AlphaFold3. B. Active site of 5NCB with guaiacol docked as per [33], in close proximity to heme co-factor (3.9 Å). C. Active site of crystal structure of SwGdmA, PDB 7OWT, with guaiacol docked in close proximity to catalytic mononuclear iron (4.8 Å). D. Active site of 7OWT with docked guaiacol and overlaid candidate demethylases. Side chains highlight residues implicated in substrate recognition (7OWT residues: I165, I207, F223). Modelled structures show similar placements of these residues, with the exception of PtGdmA and MpGdmA, which have threonine and isoleucine in place of F223, respectively. E. activity of candidate demethylases with 5 mM guaiacol. Both CnGdmAB and BaGdmAB are able to completely convert guaiacol to catechol, with the benchmark GcoAB capable of only 22% conversion. F. Temperature sensitivity of CnGdmAB and BaGdmAB during protein expression. Both candidates are able to completely demethylate 10 mM guaiacol when expressed at 18 °C, however, BaGdmAB shows greater sensitivity to increased expression temperatures, achieving only 45% conversion compared to 61% for CnGdmAB when expressed at 25 °C.

To identify candidate Rieske demethylases suitable for integration into our pathway, we performed a BLASTp search using the guaiacol demethylase from *Cupriavidus necator*, previously shown to efficiently demethylate guaiacol and 3-*O*-methylguaiacol. Homologs were filtered by sequence similarity (>75%) to both the oxygenase (*gdmA*) and its cognate reductase (*gdmB*) located at the same genomic locus. Five *gdmAB* pairs were selected for functional screening against *gcoAB*, all retaining conserved residues implicated in catalysis (Figure 2D). To maximize electron transfer capacity, the reductase (*gdmB*) was assembled upstream of the oxygenase (*gdmA*) within a single operon via JUMP assembly[28], mirroring native gene organization and ensuring preferential expression of the redox partner[29,30].

Whole-cell biotransformations in *E. coli* BL21(DE3) revealed marked differences in activity. Whereas GcoAB converted only ∼3.1 mM guaiacol (63% conversion), the Rieske demethylases from *C. necator* (CnGdmAB) and *Burkholderia anthina* (BaGdmAB) achieved complete demethylation of 5 mM guaiacol (Figure 2E). In contrast, homologs from *Paraburkholderia solitsugae* and *Methylibium petroleiphilum* showed reduced activity relative to GcoAB, while the enzyme from *Sphingobium wittichii* exhibited negligible activity. This trend mirrors prior observations in which *Cn*GdmAB, but not *Sw*GdmAB, supported growth on guaiacol as a sole carbon source when expressed in *Pseudomonas putida* KT2440, despite similar in vitro activity, suggesting expression or folding limitations in vivo[26].

Consistent with this hypothesis, SDS-PAGE analysis revealed detectable expression only for *Cn*GdmAB and BaGdmAB, although predominantly in the insoluble fraction. Biotransformations at higher substrate loadings (10 mM guaiacol) produced similar catechol titres across all constructs without reaching completion, indicating that enzyme abundance, rather than intrinsic catalytic efficiency, limited turnover under these conditions.

To select a demethylase for pathway integration, we compared expression robustness across temperatures. Although *Cn*GdmAB and *Ba*GdmAB exhibited similar catalytic activity, *Cn*GdmAB displayed greater tolerance to elevated expression temperatures (Figure 2F). Given prior links between thermotolerance and improved in vivo performance of Rieske enzymes[31], *Cn*GdmAB was selected for downstream pathway construction.

### Alternative demethylases increase substrate flux and adipic acid titres

We next examined whether the improved demethylation observed for CnGdmAB would translate to enhanced performance in the full adipic acid biosynthetic cascade. This pathway couples two oxygen-dependent reactions – guaiacol *O*-demethylation and intradiol catechol ring cleavage by catechol 1,2-dioxygenase (CatA) – with reduction of *cis,cis*-muconic acid (ccMA) by the enoate reductase from *Bacillus coagulans* (BcER). BcER sequentially reduces ccMA to 2-hexenedioic acid and then adipic acid using electrons from NADH transferred via FAD, a 4Fe-4S cluster, and FMN. Overall, the pathway requires multiple metal centres and flavin cofactors, making balanced expression and maturation particularly challenging (Fig. 3A).

**Figure 3.**
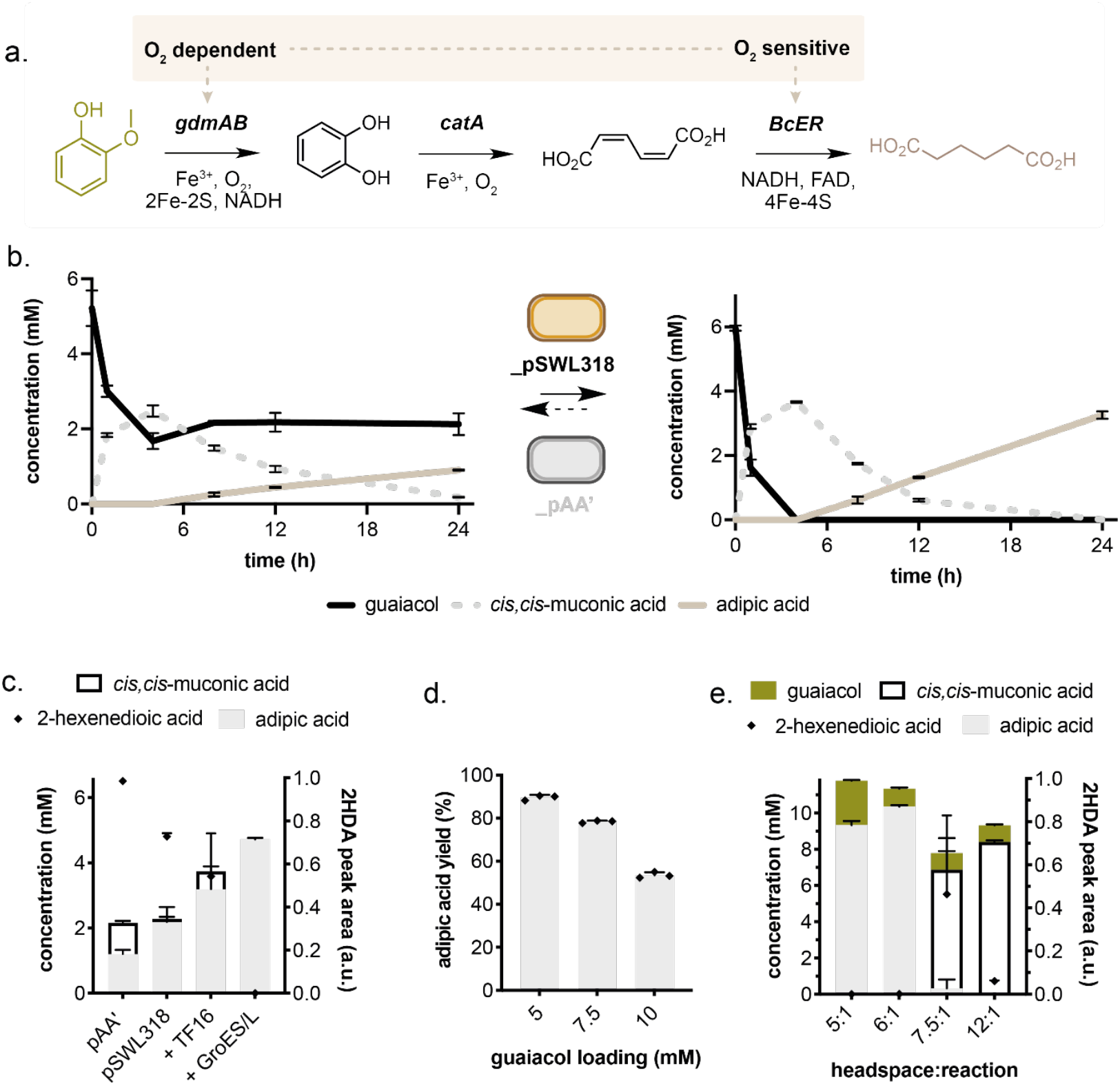
A. Full pathway expressed in E. coli for the production of adipic acid from guaiacol, highlighting oxygen dependent enzymatic steps (CnGdmAB and CatA) and the oxygen sensitive reduction of muconic acid to adipic acid by BcER. B. Time-course of full pathway production of AA using previously published pAA’*[8]* containing heme-dependent GcoAB demethylases, in comparison to pSWL318, containing CnGdmAB. pAA’ produces 1.0 mM adipic acid in 24 h, with substantial residual guaiacol, whereas pSWL547 produces 3.1 mM adipic acid within the same timeframe. C. Optimisation of protein-expression conditions to improve adipic acid production from guaiacol. Co-expression of TF16 and GroES/EL improve adipic acid at 5 mM loading. D. Guaiacol substrate loading increase reduces overall adipic acid yield. E. Increased headspace ratio dramatically reduces the activity of BcER, accumulating muconic and hexenedioic acid intermediates.

The *C. necator gdmAB* genes were introduced into the pAA plasmid backbone, which co-expresses CatA and BcER under IPTG control, generating pSWL318. This enabled direct comparison to our previously reported GcoAB-based system. When incorporated into the full pathway, the enhanced demethylation capacity of the Rieske enzymes was retained, resulting in substantially increased adipic acid titres relative to the P450-based system (Figure 3B). These results confirmed that relieving the entry-point bottleneck increases flux through the entire cascade.

### Recombinant protein limits pathway productivity

Despite improved demethylation, time-course experiments revealed incomplete conversion to adipic acid within 24 h at moderate substrate loadings. Accumulation of ccMA and 2-hexenedioic acid suggested that BcER had become a new bottleneck. We therefore focused on improving recombinant protein quality by minimizing misfolding and promoting holoenzyme formation.

To increase experimental flexibility, pathway genes were re-cloned into the modular JUMP system [32], generating plasmid pSWL337. Screening alternative plasmid backbones revealed no correlation between copy number and adipic acid production (Supplementary Figure 1), and a medium-copy pBR322 origin was selected.

Encapsulation of cells in sodium alginate, previously shown to protect BcER activity[9], proved incompatible with guaiacol demethylation, likely due to diffusion limitations (Supplementary Figure 2). In contrast, co-expression of molecular chaperones significantly improved pathway performance. Both GroES/L and trigger factor (Tf) enhanced productivity, enabling complete conversion at lower substrate loadings (Figure 3C). However, increasing guaiacol concentrations again led to reduced yields (Figure 3D), indicating additional limitations.

Given the extensive use of Fe-S clusters in the pathway, we hypothesized that incomplete cofactor incorporation impaired enzyme maturation. Although genetic restoration of SUF machinery did not improve yields (Supplementary Figure 3A, B), supplementation with Fe(II) and Fe(III) salts during expression significantly enhanced productivity, with optimal performance at ∼0.2 mM iron (Supplementary Figure 3C, D). Further gains were achieved by limiting gas transfer during protein expression, likely reducing oxidative damage to the oxygen-sensitive 4Fe-4S cluster of BcER. Under microaerobic expression conditions, adipic acid titres increased from ∼3 mM to 8.4 mM at 82% yield (Supplementary Figure 4).

### Process optimisation enables near-quantitative conversion

With improved active enzyme expression, we optimized reaction parameters. Reducing glucose concentration from 3% to 1.5% increased adipic acid titres without limiting biotransformation (Supplementary Figure 5), likely by reducing acidification resulting from fermentation.

Because upstream oxidation steps require oxygen while BcER is oxygen-sensitive, we systematically tuned oxygen availability by varying cell density and reaction headspace. Performance was robust between OD_600_ = 36-48 (Figure 6A), and simple adjustment of reaction volume substantially improved titres (Figure 3E). Under optimal conditions, 10 mM adipic acid (>95% conversion) was achieved. Higher guaiacol loadings resulted in residual substrate, and although guaiacol demethylation generates formaldehyde, co-expression of additional formaldehyde detoxification genes did not improve demethylation or product formation (Supplementary Figure 7), indicating that formaldehyde toxicity is not the primary limitation under these conditions. Instead, this likely reflects increased oxygen demand and conflicting requirements of upstream and downstream enzymes (Supplementary Figure 8).

Combining these insights, we constructed a single-plasmid system (pSWL547) encoding the full pathway and auxiliary chaperones (Figure 4A), which outperformed the two-plasmid configuration (Supplementary Figure 9). Under optimized conditions, this system achieved rapid demethylation of 10 mM guaiacol and near-complete conversion to adipic acid within 24 h (Figure 4B).

**Figure 4.**
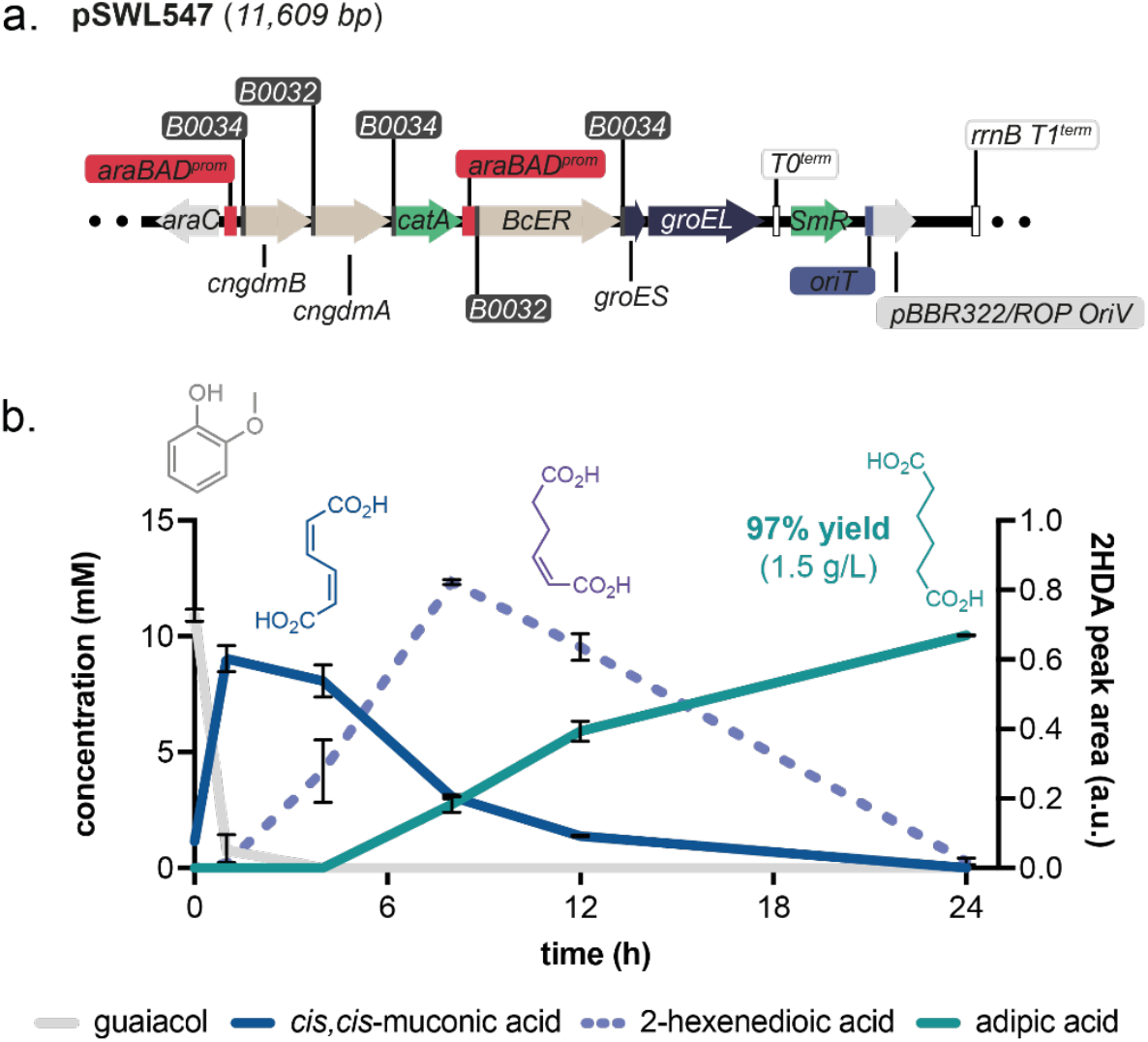
A. Plasmid map of pSWL547 for efficient conversion of guaiacol to adipic acid. B. Time-course of the biotransformation of guaiacol to adipic acid under optimised experimental conditions. Guaiacol is rapidly demethylated, with 90% reduction of guaiacol within the first hour, mirrored by a concomitant increase in *cis, cis-*muconic acid, followed by a gradual reduction to 2-hexenedioic acid and then adipic acid, achieving a final titre of 10.0 mM within 24 h (1.5 g/L, 97% yield).

Overall, the integration of improved demethylases, enhanced recombinant protein quality, and refined reaction engineering yielded a system capable of producing 1.5 g/L adipic acid at >95% yield under standard laboratory conditions.

## Discussion

While this study establishes an efficient single-strain, one-pot route for the microbial conversion of lignin-derived guaiacol to adipic acid, further gains in productivity are likely to be achieved through targeted bioprocess intensification. In particular, operation in controlled bioreactors will enable precise regulation of dissolved oxygen (dO2), a key parameter given the contrasting oxygen requirements of the pathway’s terminal enzymes. Fine-tuning oxygen availability offers a promising strategy to balance the demands of the oxygen-dependent demethylation and dioxygenation steps with the oxygen sensitivity of the downstream enoate reductase, and may unlock higher substrate loadings and product titres than are accessible in shake-flask cultures. Beyond process control, adaptive laboratory evolution (ALE) and enzyme-focused engineering represent complementary routes to further pathway optimization. While replacement of the rate-limiting guaiacol demethylation step with Rieske monooxygenases was central to the performance gains reported here, additional improvements may be achieved by identifying more active or oxygen-tolerant variants of downstream enzymes. Systematic exploration of homologous enzymes or directed evolution approaches could further enhance pathway robustness under industrially relevant conditions. Finally, comprehensive life-cycle assessment (LCA) will be essential to quantify the environmental benefits of this bioprocess relative to conventional petrochemical adipic acid synthesis, particularly as the system is extended to real, heterogeneous waste streams derived from lignin depolymerization. Together, these efforts will help define the industrial potential of microbial lignin valorization routes to commodity chemicals.

## Conclusions

We report an efficient whole-cell biocatalytic platform for converting lignin-derived guaiacol into adipic acid using engineered *Escherichia coli*. By exploiting natural enzyme diversity to replace the rate-limiting guaiacol demethylation step, we significantly improved substrate processivity and increased flux through the entire synthetic pathway, enabling near-quantitative conversion in a single strain and single vessel. Combining targeted enzyme selection with optimization of protein expression and reaction conditions led to a more than threefold improvement in performance, increasing adipic acid production from 0.45 g/L at 65% yield to 1.46 g/L at 95% yield under mild aqueous conditions. Beyond performance gains, this pathway integrates radical monooxygenase chemistry, enzymatic aromatic ring cleavage, and iron-sulfur-dependent alkene reduction, a combination of transformations not readily accessible through existing synthetic methods, highlighting the power of engineered microbes to access complex chemical space while enabling sustainable routes to commodity chemicals.

