## Supplementary Information for "Heterologous Rieske non-heme iron monooxygenases enable efficient microbial conversion of lignin guaiacol to adipic acid"

### S1. General materials and methods

Unless otherwise stated, starting materials and reagents were obtained from Sigma and used without further purification. All water used experimentally was purified with a Suez Select purification system (18 MΩ.cm, 0.2 μM filter).

The following abbreviations are used throughout: adipic acid (AA), cis,cis muconic acid (ccMA), 2-hexenedioic acid (2-HDA).

#### S1.1. Identifying candidate demethylases

A protein BLAST search[1] was conducted on both CnGdmA and CnGdmB, for which candidate pairs were selected based on the following criteria: hits for GdmA >75%, the corresponding GdmB pair was identified in the top one hundred hits, and both genes were found within the same locus in the host genome. These genes were synthesised by IDT and expressed in *E. coli* BL21 (DE3).

#### S1.2. Molecular biology

PCR reactions were carried out using Phusion High-Fidelity DNA Polymerase (NEB), following the manufacturer recommendations. SapphireAmp 2x Master Mix (Takara Bio) was used for colony PCR reactions. Oligonucleotide primers were designed in silico and synthesized by Integrated DNA Technologies. PCR reactions were checked by agarose gel electrophoresis and final plasmids confirmed by Sanger sequencing (Azenta). Restriction enzymes and T4 DNA ligase were purchased from Thermo Fisher Scientific or NEB.

Cloning of the pQLinkN expression plasmids were assembled as has been described previously[2,3] using iterative integration of BcER, CatA and the respective demethylases (GdmAB or GcoAB). 5' BamHI and 3' NotI recognition sites were added by PCR and followed by digestion-ligation.

Cloning of the pJUMP vectors was performed as in Valenzuela-Ortega and French[4]. Gene sequences were amplified with appropriate overhangs (AGCC and TTCG for O parts, AATG and GCTT for NOC parts). NOC parts are gene-encoding sequences which cover the entirety of the gene, including the start and stop codons. O parts are similarly gene-encoding sequences, however they do not contain the start of stop codons, instead these are supplied by the corresponding RN (ribosome-binding site with associated start codon) as well as a corresponding CT part (RNA polymerase terminator sequence with corresponding stop codon) by PCR followed by Goldengate assemblies[5] with BsaI/BsmBI and T4 DNA ligase (New England Biolabs). Assemblies for demethylases were performed to produce operonic units in a single assembly with *GdmB* was designed as an NOC part followed by *GdmA* as an O part.

2-plasmid screening with chaperones involved assembly of *CnGdmBA*, *CatA* and *BcER* assembled on a single kanamycin resistant plasmid (pJUMP29) with cells co-transformed with pGro7 (Takara)[6]. For single-plasmid expression, the GroES/EL region of pGro7 was amplified by PCR to introduce BsaI recognition sites to generate a single NOC part, which was assembled as a JUMP part under the control of PBad promoter. This chaperone part was then assembled with *CnGdmBA*, *CatA*, and *BcER* into a pJUMP49 backbone.

#### S1.3. High Performance Liquid Chromatography

Biotransformation samples were quenched with an aqueous acid solution of 55mM HCl, 0.1% TFA with 0.01g/L caffeine as an internal standard, in a 1:3 sample-to-quencher ratio. Prior to HPLC analysis, samples were boiled for 1 hour, cooled to room temperature and centrifuged at 13 000 *rcf* for 15 minutes to remove cell debris. Analysis was performed at a flow rate of 0.4 mL/min, 30 °C, 10 µL sample injection. The following mobile phases were used; miliQ $H_2O$ + 0.1% trifluoroacetic acid (TFA), and acetonitrile + 0.1% TFA. The mobile phase gradients are depicted in Table S1.

*Table 1 Mobile phase gradients for HPLC*

| Time (min) | MiliQ $H_2O$ + 0.1% | Acetonitrile + 0.1% TFA |
|---|---|---|
| 0 - 5 | 95 | 5 |
| 5 - 10 | 85 | 15 |
| 10 - 20 | 50 | 50 |
| 20 – 20.2 | 95 | 5 |
| 20.2 to 25 | 95 | 5 |

HPLC analysis was performed using a Thermo Scientific Dionex UltiMate 3000 UHPLC with UV detector reading at 206nm using a HyperSil Gold18 column (150 x 3 mm x 3um, ThermoFisher).

### S2. Strains, media and culturing conditions

#### S4. 1 Strains

Routine cloning was carried out with *E. coli* DH5α, and biotransformations performed predominantly in *E. coli* BL21 (DE3). Investigations into effects of SUF operon loss were carried out in MG1655 (Wild-type comparator) or BL21 (DE3) complemented with the sufABCDSE or SufCDSUB operon[7] for complementation.

#### S4. 2 Media

Media used for bacterial growth was either lysogeny broth (LB; 10 g/L bacto-tryptone, 5 g/L yeast extract, 10 g/L NaCL) or Terrific broth (TB; 24 g/L yeast extract, 20 g/L tryptone, 0.4% glycerol, 12.5 g/L K2HPO4).

Reactions were carried in M9 minimal media: $Na_2HPO_4$, (6 g/L), $KH_2PO_4$ (3 g/L), $NH_4Cl$ (1 g/L), NaCl (0.5 g/L), $MgSO_4$ (241 mg/L) and $CaCl_2$ (11 mg/L) with an appropriate amount of starting substrate and glucose.

#### S4. 3 Culturing conditions

5mL of overnight cultures was used to inoculate 250 mL fresh media in 500 mL GL 45 thread Duran® baffled flasks (DWK Life Sciences) and grown at 21-30 ℃ with 220 rpm shaking until $OD_{600}$ 0.4-0.6. Once target $OD_{600}$ was achieved, protein expression was induced by addition of 0.4% arabinose, with 0.4mM IPTG included for chaperone co-expression and SUF overexpression screens. Additional additives including ammonium iron(III) citrate (0.76mM unless otherwise stated), iron(II) sulfate (0.1-1 mM) and 5-aminolevulinic acid (0.76 mM, *GcoAB* expressing plasmids only) were added at the point of induction. Post-induction cultures were tightly sealed and incubated at 21 ℃ for 16-20 h for protein expression.

For the expression of the demethylase homologs, cultures were grown to $OD_{600}$ ~0.6 at 25 °C in 250 mL LB in a 500 mL baffled flask, cultures were induced with 0.4% L-arabinose and 0.5 mM ammonium Fe(III) citrate was added. Cultures were left to express at 16 °C, 220 rpm for 20 h before harvesting for whole cell biotransformations. 1 mL of culture was taken for SDS-PAGE analysis to compare with an empty vector control.

### S3. Whole-cell biotransformations

Cultures were harvested by centrifugation at 2700 x rcf at 10 °C for 10 mins. The supernatant was decanted and pellets resuspended with PBS. The resuspension was once more centrifuged at 2700 x rcf, 10 °C, 10 mins. Reactions were set up by resuspending cells in M9 minimal media with the desired amount of substrate to the target $OD_{600}$. Unless otherwise stated, reactions were carried out in a 15 mL screw-cap polypropylene tube (120 x 17 mm) with 3 mL of resuspended cells in the reaction buffer. The reactions were left to incubate for 24 h at 21 °C, 220 rpm for 24 h before quenching.

### S4. Supplementary figures

#### S4. 1 Predicted plasmid copy number does not correlate with in vivo pathway performance

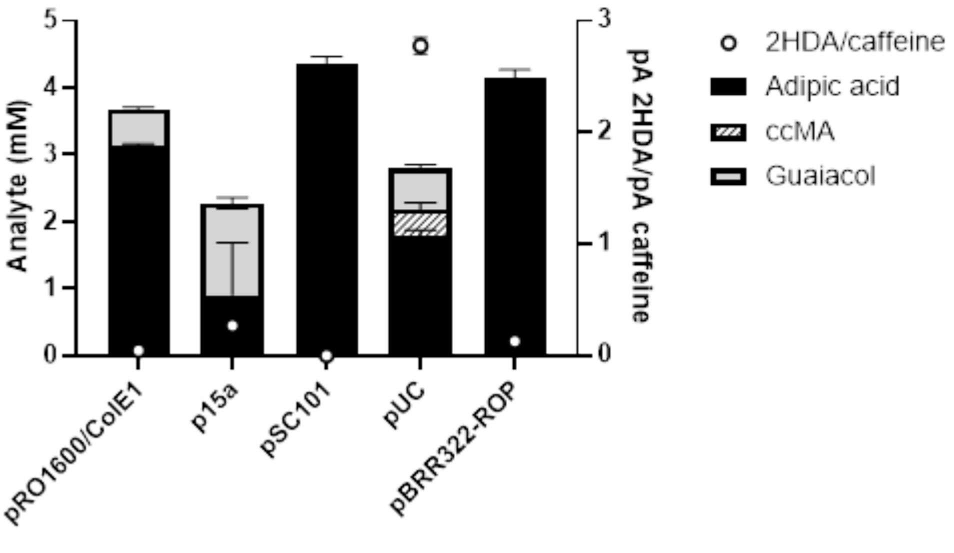

Figure S1. Screening different copy number plasmid backbones showS no correlation between predicted copy number and performance of the heterologous guaiacol to adipic acid pathway.

#### S4. 2 Cell immobilisation with alginate impairs demethylation of guaiacol to catechol and reduces product titres

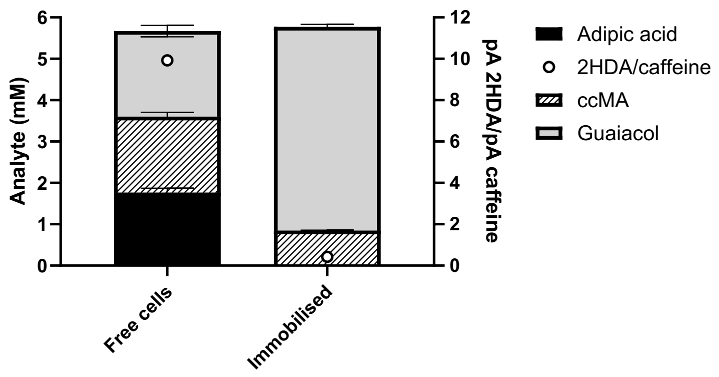

Figure S3. Cells expressing guaiacol to adipic acid pathway enzymes either resuspended directly in reaction buffer, or first immobilised in alginate beads. Elimination of adipic acid production and accumulated residual guaiacol under immobilised conditions suggests poor diffusion of guaiacol through alginate beads.

#### S4. 3 Effects of Fe-S maturation genes and Fe(II/III) supplementaition on biotransformation efficacy

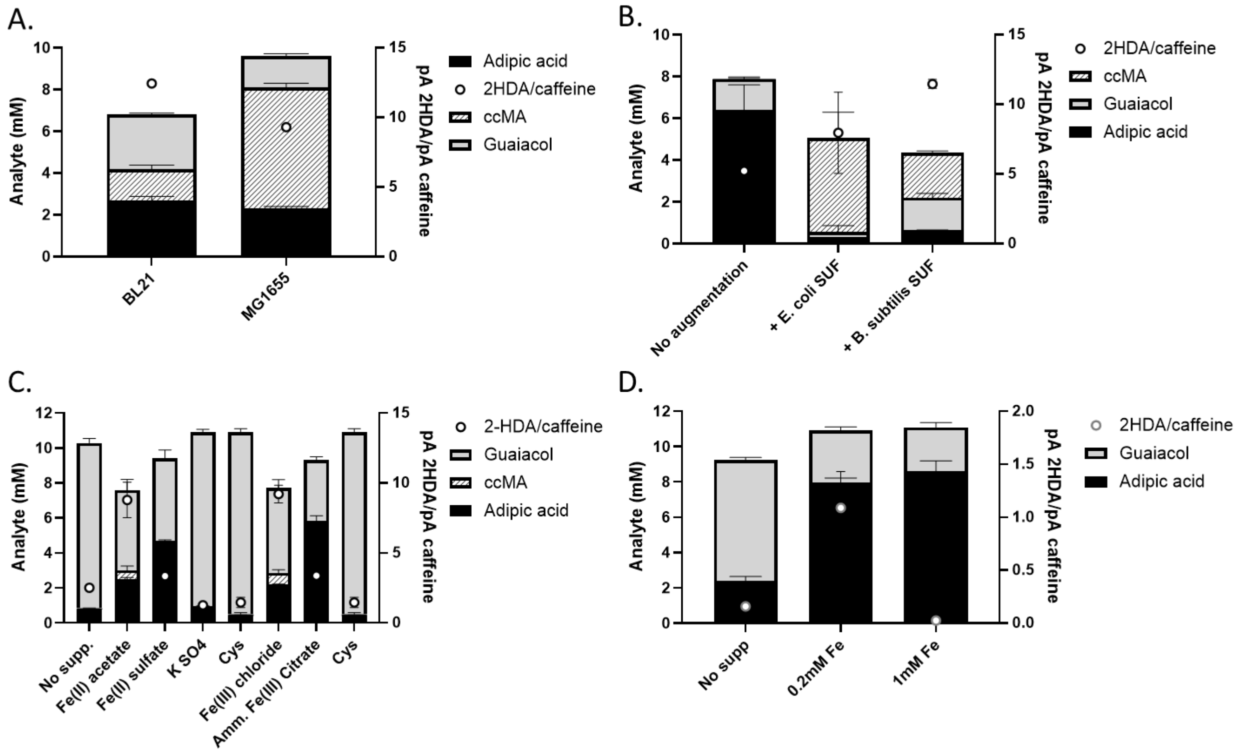

Figure S3. A) comparison of MG1655 (functional SUF operon) and BL21 (defective SUF operon) shows significant accumulation of ccMA despite the functional Fe-S maturation pathways in wildtype *E. coli*.

B) complementation of BL21 with either the SUF operon from *E. coli* MG1655 or Bacillus subtilis has a negative impact on ene-reductase activity leading to both reduced adipic acid titres and increased ccMA accumulation. C) Screening iron and sulfur salts identifies Fe(II) $SO_4$ and Fe(III) citrate as additives to improve titres of adipic acid at higher substrate loadings. D) Improvements to product titres from combined Fe(II) and Fe(III) supplementation rapidly plateau after 0.2 mM addition.

#### S4. 4 Controlling aeration during protein expression significantly enhances adipic acid titres

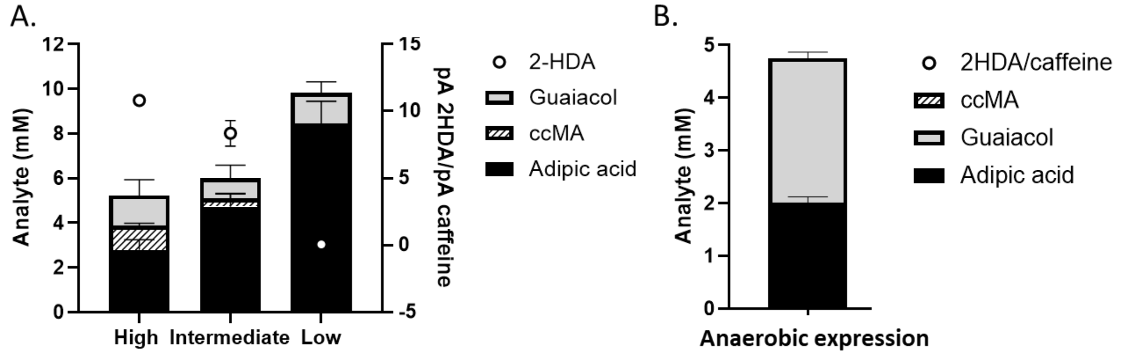

Figure S4. A) Generating a micro-aerobic environment by limiting gas exchange during protein expression, followed by aerobic biotransformations, improves adipic acid titres from ~3.1 mM to >8 mM. B) Anaerobic protein expression followed by aerobic biotransformations impairs guaiacol *o*-demethylation. Substantial residual guaiacol indicates poor demethylase activity, however a lack of quantifiable ccMA and 2-HDA suggests production of a highly active enoate reductase.

#### S4. 5 Effect of carbon exogenous carbon source during biotransformations

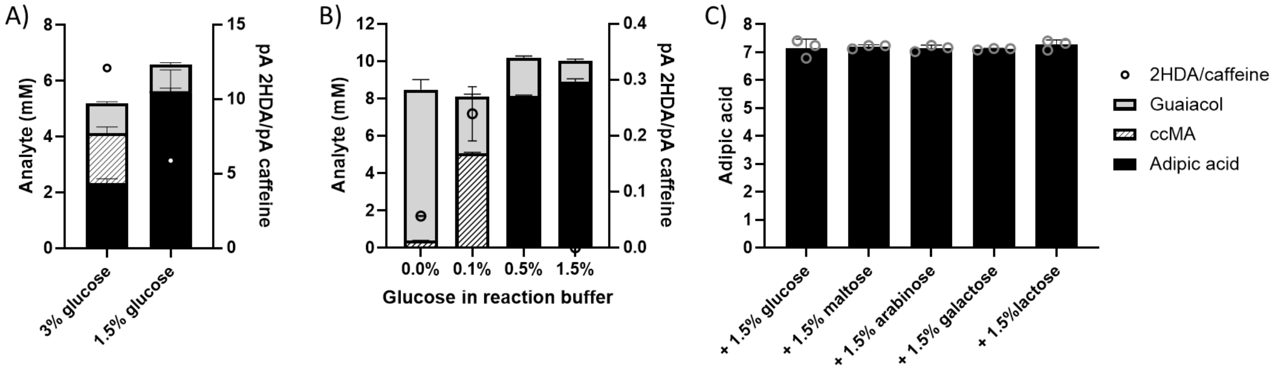

Figure S6. Supplementing reaction buffer different combinations of carbon sources to generate reducing equivalents for biotransformations. A) reducing glucose supply from 3 % to 1.5 % more than doubles adipic acid titres from 2.3 mM to 5.6 mM. B) The glucose concentration in reaction buffers can be reduced to 0.5% with minimal impact on product titres, however further decreases severely limit catalytic activity. C) Supplementing 1.5 % glucose with 1.5 % (w/v) additional carbon sources shows no impact on adipic acid titres suggesting that at 1.5% glucose the availability of carbon source is not limiting further improvements to adipic acid synthesis.

#### S4. 6 Increasing oxygen availability during biotransformation improves demethylation at the expense of reductase activity

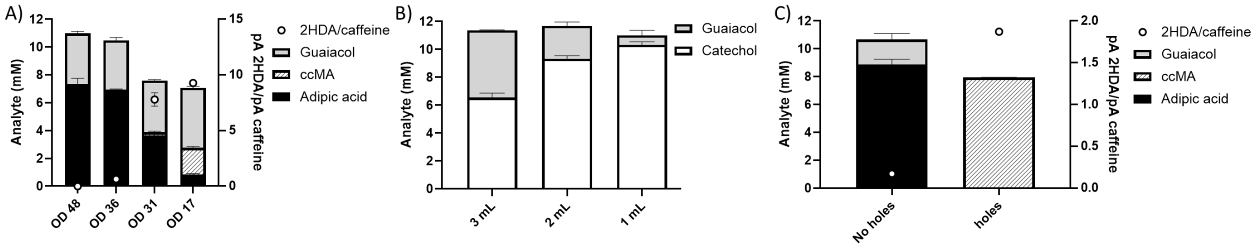

Figure S6. Effect oxygen availability during biotransformations. A) reducing cell density from $OD_{600}$ = 48 to $OD_{600}$ = 36 shows negligible impact on adipic acid titres. Further decreases result in rapid decreases in adipic acid formation and increased 2-HDA and ccMA accumulation. B) Increasing reaction headspace by decreasing reaction volumes improves guaiacol demethylation. C) Increasing oxygen availability by introducing holes into reaction containers allows complete demethylation of 10 mM guaiacol, but abolishes enoate reductase activity.

#### S4.7 Formaldehyde detoxification does not improve guaiacol demethylation

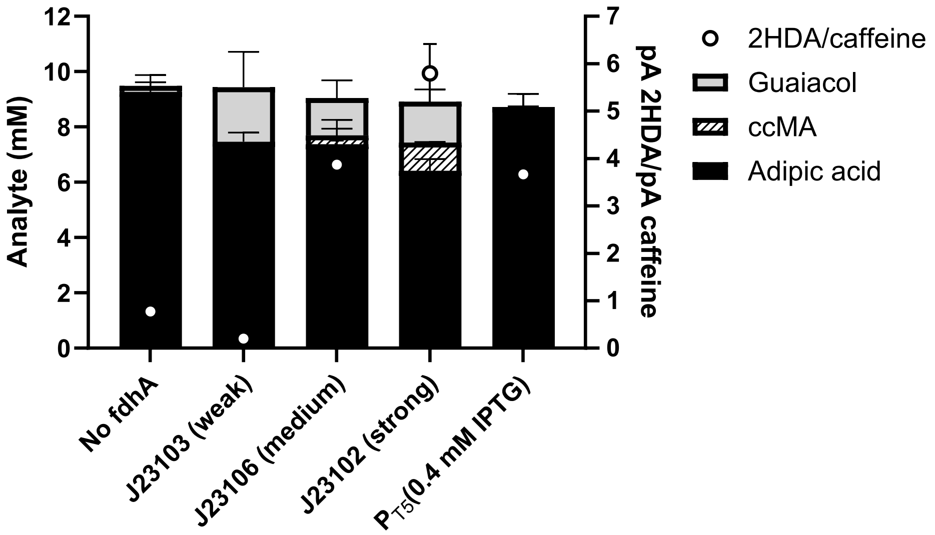

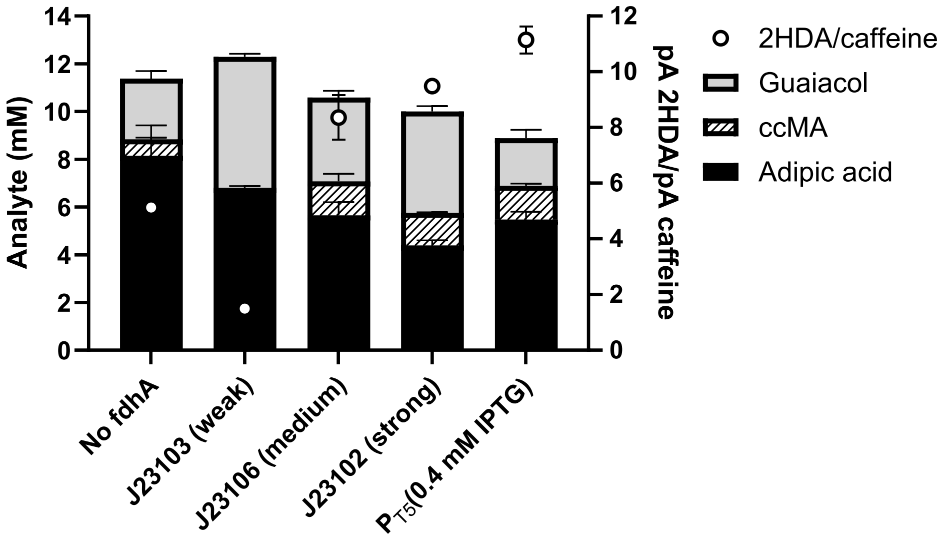

Figure S7. Complementation of strains with formaldehyde dehydrogenase from *Pseudomonas putida*(*fdhA*) does not improve guaiacol demethylation and reduces adipic acid titres. Biotransformation strains were complemented with plasmids expressing *fdhA* under weak (J23103), medium (J23106), strong (J23102) constitutive promoters, or under the control of the IPTG-inducible $P_{T5}$ promoter. Top panel shows biotransformations with 10 mM and lower panel with 12 mM guaiacol loading. In all cases neither guaiacol demethylation nor final product titres were improved by the presence of *fdhA* expression.

#### S4. 8 Reducing enzymatic oxygen requirements can allow higher adipic acid titres

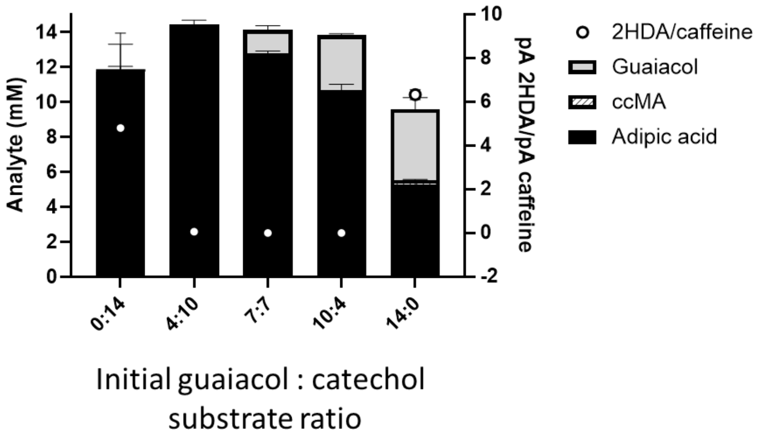

Figure S8. Effects of changing substrate composition on biotransformation efficiency. Reactions with lower oxygen-dependent demethylation requirements were able to achieve higher titres of adipic acid than those with both oxygen-dependent demethylation and oxidative intradiol ring cleavage.

#### S4. 9 Enhancement of activity by plasmid consolidation

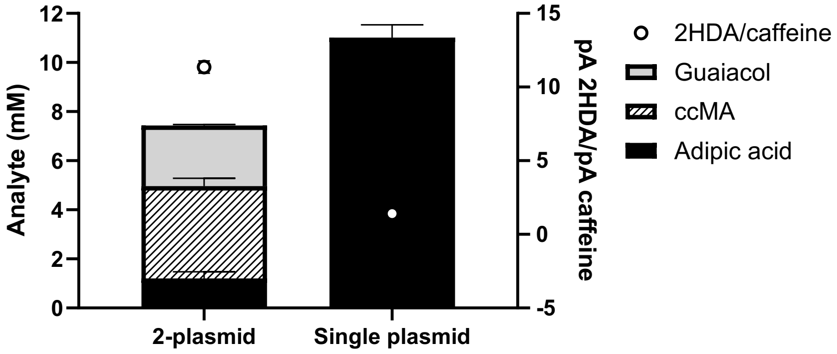

Figure S9. Combining chaperones GroES/L onto the same plasmid as the guaiacol to adipic acid pathway genes significantly improves pathway performance at high guaiacol loadings.

### S5. Table of genes

| Gene | Nucleotide sequence (5’ to 3’) |
|---|---|
| *gcoA* | ATGACCACCACCGAACGTCCGGATCTGGCATGGCTGGATGAAGTTACCATGACACAG<br>CTGGAACGTAATCCGTATGAAGTTTATGAACGTCTGCGTGCAGAAGCACCGCTGGCAT<br>TTGTTCCGGTTCTGGGTAGCTATGTTGCAAGCACAGCCGAAGTTTGTCGTGAAGTTGCA<br>ACCAGTCCGGATTTTGAAGCAGTTATTACACCGGCAGGCGGTCGTACCTTTGGTCATC<br>CGGCAATTATTGGTGTTAATGGTGATATTCATGCCGATCTGCGTAGCATGGTTGAACCG<br>GCACTGCAGCCTGCAGAAGTTGATCGTTGGATTGATGATCTGGTTCGTCCGATTGCAC<br>GTCGTTATCTGGAACGTTTTGAAAATGATGGTCATGCAGAACTGGTTGCCCAGTATTGT<br>GAACCGGTTAGCGTTCGTAGCCTGGGTGATCTGCTGGGTCTGCAAGAAGTGGATAGC<br>GATAAACTGCGTGAATGGTTTGCAAAACTGAATCGCAGCTTTACCAATGCAGCAGTTGA<br>TGAAAATGGCGAATTTGCCAATCCGGAAGGTTTTGCCGAAGGTGATCAGGCAAAAGCA<br>GAAATTCGTGCAGTTGTTGATCCGCTGATTGATAAATGGATTGAACATCCTGATGATAG<br>CGCAATTAGCCATTGGCTGCATGATGGTATGCCTCCGGGTCAGACCCGTGATCGTGA<br>ATATATCTATCCGACCATTTATGTTTACCTGCTGGGTGCAATGCAAGAACCTGGTCATG<br>GTATGGCAAGCACCCTGGTTGGTCTGTTTAGCCGTCCGGAACAACTGGAAGAGGTTGT<br>TGATGATCCGACACTGATTCCGCGTGCCATTGCAGAAGGTCTGCGTTGGACCTCACC<br>GATTTGGAGCGCAACCGCACGTATTAGCACCAAACCGGTTACCATTGCCGGTGTTGAT<br>CTGCCTGCAGGTACACCGGTTATGCTGAGCTATGGTAGCGCAAATCATGATACCGGTA<br>AATATGAAGCACCGAGCCAGTATGATCTGCATCGTCCGCCTCTGCCGCATCTGGCCTT<br>TGGTGCAGGTAATCATGCATGTGCAGGTATCTATTTTGCGAATCATGTTATGCGTATTGC<br>CCTGGAAGAACTGTTTGAAGCAATTCCGAACCTGGAACGTGATACCCGTGAAGGTGTT<br>GAATTTTGGGGTTGGGGTTTTCGTGGTCCGACCAGCCTGCATGTGACCTGGGAAGTTT<br>GA |
| *gcoB* | ATGACCTTTGCAGTTAGCGTTGGTGGTCGTCGTGTTGATTGTGAACCGGGTCAGACCC<br>TGCTGGAAGCATTTCTGCGTGGTGGTGTTTGGATGCCGAATAGCTGTAATCAGGGCAC<br>CTGTGGTACATGTAAACTGCAGGTTCTGAGCGGTGAAGTTGATCATGGTGGTGCACCG<br>GAAGATACCCTGAGCGCAGAAGAACGCGCAAGCGGTCTGGCACTGGCATGTCAGGC<br>ACGTCCGCTGGCAGATACCGAAGTTCGTAGCACCGCAGATGCAGGTCGTGTTACCCA<br>TCCGCTGCGTGATCTGACCGCAACCGTTCTGGAAGTTGCAGATATTGCACGTGATACC<br>CGTCGTGTTCTGCTGGGTTTAGCAGAACCGCTGGCATTTGAAGCAGGTCAGTATGTTG<br>AACTGGTTGTTCCTGGTAGCGGTGCACGTCGTCAGTATAGCCTGGCAAATACAGCAGA<br>TGAAGATAAAGTTCTGGAACTGCATGTTCGTCGTGTGCCAGGTGGTGTTGCAACCGAT<br>GGTTGGCTGTTTGATGGTCTGGCAGCCGGTGATCGTGTTGAAGCAACCGGTCCGCTG<br>GGTGATTTTCATCTGCCTCCGCCTGATGAAGATGATGGTGGTCCGATGGTTCTGATTGG<br>TGGTGGTACAGGCCTGGCACCGCTGGTTGGTATTGCCCGTACCGCACTGGCACGCC<br>ATCCGAGCCGTGAAGTGCTGCTGTATCATGGTGTTCGTGGTGCAGCAGATCTGTATGA<br>TCTGGGTCGTTTTGCAGAAATTGCCGAAGAACATCCGGGTTTTCGTTTTGTTCCGGTTCT<br>GAGTGATGAACCGGATCCTGCATATCGTGGTGGTTTTCCGACCGATGCATTTGTTGAAG<br>ATGTTCCGAGCGGTCGTGGTTGGAGCGGCTGGCTGTGTGGTCCGCCTGCAATGGTTG<br>AAGCCGGTGTTAAAGCATTTAAACGTCGTCGTATGAGTCCGCGTCGTATTCATCGTGAA<br>AAATTCACACCGGCAAGCTGA |
| *cnGdmA* | ATGGTAGCCAAAACCCAAAGTCCTTATCTGATGAATGCCTGGTACGTCGCCGCGCTGT<br>CGACGGAAGTGGGACCTGAGGAGCTGTTTCATAGAAAAATCCTGGACGTGTCAGTGAT<br>GATTTATCGTAAACAGGACGATGGCACCCCTGTCGCGATGCATGATCGGTGTCCACAT |

| | |
|---|---|
| | CGATTTGCTCCGCTGCATTTAGGCAAACGGCACGGTGACGAGGTTTCATGCCCGTATC<br>ACGCGCTTCGCTTCGATTGTTTGGGGACGTGCACACACAATCCACACGGGACCGGCC<br>ATATTCCGAAAGCCGCAAGTGTGAAAACCTTTCCGCTCGCCGAAAAGTACGGTTTCGT<br>TTGGATATGGATGGGTGATGAGGCGCCAGATTTAAAGCTGCTGCCGGATTACAGCGAA<br>TTAGATATTGGTCACGCGAACGCTGTAGGCTACACGTATATGCACATGCCGGTAAACT<br>ATGAACTGATTATCGATAACGTGATGGACCTCAGCCACATTGACCATGTGCATGGTGAA<br>ATCATAACAACTCGTGGCAAGCTGTCCCCGGTCATTCCGAAAGTGGCAGATGTTCAGC<br>GCACCGTGAATGCCAGGTGGGAATGGACACAGAGTCCGGCGATGATGATCTTTGCAA<br>ACTTTTTGCCTCAGCCAGAAGAGGAAGCGCGCCACTATTTCGATATTACGTGGGCACC<br>GCCCGCCAACATTCAGCTTTCTGTAGGAGCCGTTCAGGGTGAAATGTCGTTTGATGATT<br>ACATCGGCCAATATGATTTGCATACCACCACCCCCGAAACTCAAGGCAGCACGCATT<br>ACTTTTTCGCGACTCGTCGTAATCATATCGTTGAAGACGGCGATTATAACAAAATGAAA<br>ATTCAGGCCATGCATGGGGCGTTCGAAAATGAAGACGGACCCATCATTTCCGCGGTA<br>CAGCGTGAAATGGGTGACGCAAACTTTTTCGATCTGAATCCGGTTCTAATGTCTAATGA<br>CGTCGCAGCTGTCAAAGTGCGCCGCAAGCTGGAGATGCTTATCAGCGCTGAACGCC<br>AACGTAAACTGGAGGATTCGTGA |
| *cnGdmB* | ATGAACACGCAAGATATTCATGTAGCAACCACGTTAAAAGTGTTGGTCAGGGAGAAAG<br>AAACAGTTGCCGACGGAATAGTGCGTTTCGAACTCCGCCCGACCTCTGGGGAACTGC<br>TGCCGTCATTTACCGCAGGGTCGCATATTGACGTTACCTTACCCAACGGCGCGGTGC<br>GTCAATATAGTTTATGTAATGCGCCGGACGATAACGGCCGCTACGAAATTGCGGTACT<br>GCTTGAACCGCTAGGCCGTGGTGGCTCACGTAGCGCCCATGTTGACCTGCATCGAGG<br>TTCCGTTCTGCAAATAAGCCCGCCGCGCAATCTGTTTCCACTGAGTGATGCCGGGCA<br>CTCGATCCTCATTGCAGGCGGAATTGGTATTACGCCAATTCTGAGCATGGCGCATCAT<br>CTGGCATGTGGCGGCCGGAGTTTTGAATTGCACTATTGCACACGATCTGTGGCTCGCA<br>CCGCGTTTCTTGAGCGTATCACTTCGTCCGAATTCGCGGATAGCGTCACTATCTACCAT<br>GATGATTATACTCAGCAGGAACGGTTCTCAGCAAGAAATACGCTCGCGAGTCCGACTG<br>ACGATACGCATATCTACGTATGTGGACCGACCGGTTTTATGGATCACGTTATCGGCGT<br>GGCTCTGGCTGCCGGTTGGCAGACGGCGAACGTGCACCGCGAGTATTTCGTTGCTCC<br>TAGCGCAGATACATCCGGCGATCGTCCGTTTACGGTTGAGTTCGCTCGTACCGGGCA<br>AGCCGTCGTCGTGCCACCTTCCCAGAGCGTAGCGCAGGCCCTTGCAGCGCACGGTA<br>TTGATATCCCAGTCAGCTGCGAACAGGGTATTTGCGGTACCTGCATGATGCGCGTGCT<br>GGAAGGCGAGCCTGACCATCGCGACACCTTTCTGACCACTCAGGAACACGCGGCCA<br>ATGATCGCTTTACCCCCTGCTGTTCTCGGTCGAAGACAGCCCGTTTGGTGATCGATTTC<br>TAA |
| *swGdmA* | ATGGTAGCCGGCTCTACGTTCGAGTGCCTGAAGCGGGCCTGGTACGTTGCGGCGATG<br>AGCTCGGAAGTTGAAGGGGAAGCACTGTTTCACCGTCGCATACTGGGCACTAGCGTG<br>ATGATTTATCGTCTTGCGGATGGCACCCCGGTGGCGATGCACGACCGATGTCCGCAT<br>CGTTTTGCCCCGCTGCACCTGGGGCAGCGTGAGGGAGATGAAATCGCGTGCCGTTAT<br>CATGCGCTCCGCTTTGATGCGGATGGCCGTTGCACGCACAACCCGCACGGTAACGG<br>TCGCATTCCAGACGCAGCTAGAGTACGGCGCTTTCCGTTGCTGGAACGGTATGGCTTT<br>CTTTGGATTTGGATGGATGACAGTGATCCGGATCCCGCATTACTGCCAGACTTCTCAC<br>CTCTGGAAGAGGGTCATCCTAATGCGGTCGCTCAAACCTATATGCATATGGATGTTAAC<br>TACGAGCTGATCATTGACAATGTGATGGACTTATCTCACATTGATCACGTTCATGGGGA<br>GATTATCACAACCCGTGGCCAGCTAAGTCCGGTCGTTCCTAAAGTGCGCGAGAGGGA<br>CACCCACATCAGCGCACGCTGGGAATGGTCGCAGACTCCGGCCATGATGATATTTGC<br>CCCGTTCCTGCCGCGTCCTGAGGATGCCGCACGCCATTATTTCGATATCAGTTGGACT<br>GCCCCCGCCAATATTCAGTTGTCAGTCGGCGCGGTACAAGACAGCGATGATTTTGGA<br>GATGCTACATCCCAGTACGATTTGCATACGTGTACCCCAGAAGATGCCTTTAAAACGC |

| | |
|---|---|
| | ATTACTTTTTCGCGACGCGCCGCAACCATATCGTAGATGACGCTGACTATAACCGCAA<br>AAAAATCGAAGCCATGCATGCAGCATTCGAAACCGAAGATGGTCCGATCATTACCGC<br>CGTCCAAGAAGAAATGGGTGAGGCAGAATTCTTTTCCCTCGATCCAGTGTTAATGTCGA<br>ATGATGTTGCGCCCGTGAAAGTGCGAAGACGTCTGCGTCGCATGATTGTGGAAGAAC<br>AGGCAGCTGCTGCGGCCGCGGACGCGGCTTCTTCGTGA |
| *swGdmB* | ATGAGTGGTGATTGGATTTCCGTTCGCCTGGCCGGGAAACGCCTTGTCGCGGAAGATA<br>TCGCCGCTCTTCTGCTGGAACCGGCGGATGGAACCGCGCTGCCGCCATTTGCTGCA<br>GGCGCTCACGTAGATGTGGAATTGCCGGGCGGCTTGGTCCGGCAGTATTCTCTGTGT<br>AACGCACCGGGTGACCCTGGTCGTTATGAATTAGGGGTGCTGTTGGATCGCCAGGGT<br>CGCGGTGGGAGTCGCGCGGTGCATGAGCGGCTGGCCGCTGGCCAGGCAATTCGTAT<br>TAGCCGTCCACGGAACCTGTTCCCGCTCGTCCCTGCCGCGCATTCCGTTCTGATGGC<br>CGGCGGCATCGGGATCACGCCCATCCTCGCGATGGCCGAACAACTGGCCGCAGAT<br>CGCGCGTCGTTCGAGTTACATTACTTTGTAAGATCACCAGAGCGCGCAGCGTTTGTGG<br>ATCGTCTGAGCGCTCCGAGGTTCGCAGACCGCTGTCGTCTGCACGTTGGCGATCGCA<br>TTCCGCCCTCGTTCGATGCCCCTGCCCTTTTCGACGCACCGCGCGCGGGCCACCAT<br>CTATACGTTTGCGGGCCCAATGCGTTTATGGACGCGGTGCTGGACGGTGCCAGAGCT<br>GGAGGTTGGCCGGACGATCGTCTGCATAGTGAACGATTTGCCGCGGCTCCGCTGCCA<br>GCGGGCGGTGATCGCCCGTTTGAGATCGAAGTAGAAGGTCACGGCGCAATTGTGGC<br>GGTGGCCGCAGGCCAGTCGGCTGCGGAAGCCTTGGCGGCAGCAGGTATACGTATCC<br>CTCTCTCATGCGAGCAGGGCATTTGTGGAACCTGCTTAACTACGGTGGTTGGTGGAGT<br>TCCGGACCATCGAGACAGCTATCTTAGCGATGCGGAACGTGCGGCCAATGATTGTTTT<br>ACACCGTGCTGCTCTCGTGCAGCGACCCCACGTCTGGTCATTCAATTATAA |
| *baGdmA* | ATGGTAGCCGAAACGCCTTCCCGCTATCTCATGAATGCTTGGTATGCAGCAGCGCTGA<br>GTTCTGAAGTCGGGCCGGAGGCCCTGTTCCATCGCAAGATCCTTGATGTATCGGTCAT<br>GATTTACCGTAAGCAAGACGGTACTCCAATCGCAATGCACGATCGTTGCCCGCATCG<br>CTTTGCCCCATTACACCTGGGTAAACGTCACGGTGATGAAGTGGCTTGTCCATATCAT<br>GCTCTTCGTTTTGATTGCTCCGGCGCATGCACGCATAACCCACATGGTACAGGGCATA<br>TCCCTAAAGCCGCGATCGTGCGGACGTTCCCATTGCTTGAAAAGTACGGCTTCTTGTG<br>GATTTGGATGGGGGATGCGGCCCCAGACCCGGCTCTGCTGCCAGATTACGGTGCATT<br>AGATGATGGTCATCCAAATGCTGTTGGCTATACTTACATGCACATGCCTGTGAACTACG<br>AACTTATTATTGACAATGTAATGGATTTGAGTCACATTGACCATGTACATGGCGAAATCA<br>TCACCACTCGCGGTCAGTTGTCGCCAATCGTCCCTAAAGTCACCGATGGGCAACGTG<br>CCGTTAACGCACGCTGGGAATGGACGCAGAAGCCTGCAATGATGATTTTTGCGAATTT<br>CCTCTCAAAGCCTGACGACGAGGCTCGTCATTACTTCGACATCACATGGGCTCCACC<br>AGCAAATATCCAACTTTCCGTAGGCGCTGTGCAAGGCGATAAGTCTTTCGATGAGTGTA<br>TTGGTCAGTATGATCTTCACACAACAACCCCTGAAACGTTATACTCTACTCATTACTTCT<br>TTGCAACTCGTCGGAACCATGTCGTGGAAGACGGTGACTACAATGCGATGAAAATTAA<br>AGCGATGCACGGGGCGTTTGAAAACGAGGATGGGCCTATTATCACAGCAGTACAACG<br>TGAAATGGGCGAGGCAAGTTTTTTTGACCTGGATCCGGTTCTCATGTCCAATGACGTCG<br>CCCCAGTCAAGGTACGCCGTAAGCTGGGCACACTTATTGACGCAGAACGGCGCAGC<br>GGCCAGGCAGATTCGTGA |
| *baGdmB* | ATGAACATGCAAGATATCCGCGCGGCAACAACGCTCACCGTTCAAGTCTCTGAGCGG<br>GAGCCAGTCGCAGACGGCATCGTACGGTTTGAACTTCGTCCGACATCACTGGATCCT<br>CTGCCGCCATTCACGGCGGGCGCTCATATCGATATTAAGCTCCCGAATGGTGACGTA<br>CGCCAATACAGCCTGTGTAATGCTCCTTACGTTAATGATCGCTACGAGATCGCGATCTT<br>GTTAGAGCCACAGGGCCGTGGCGGTTCGCGCTGCGCACATGCCGAATTGCAGCGCG<br>GTAGTGTGGTTCAGATTTCCGCGCCGCGCAATCTTTTCCCGCTCTCCGATGCACGCCA |

| | |
|---|---|
| | TTCCATCTTAGTGGCAGGTGGTATCGGTATCACACCGATTTTATCTATGGCGTATCAGC<br>TGGCACGCGATGGGCGCAGTTTCGACCTTCATTACTGTACCCGCAGTGCTGCTCGGG<br>CTGCTTTCCTGGAGCGTATTCACGCCTCGGAGTTTGCAGAAAATGCGGTCGTCTATCA<br>CGACGATCGGCCAGAGCATGAGCGCTTTGCCGCGAAGCATGTTTTGGCAAATCCAGC<br>AGACGACACTCACATTTATGTTTGTGGGCCAACAGGTTTCATGGATTATGTCATCGCGA<br>CCGCATTCGATGCAGGCTGGCGGGCGACCCACATTCACCGCGAGTATTTCGCCGCG<br>GCCCGCACTGAAACCCCAGGCGACTGCTCATTCACCGTCGAGTTTGCACCGACCGG<br>TCTTTCCGTTGTCGTGCCACCTTCGCAGTCGGTAGTGGAGGCATTAGCCGCTCATGGC<br>GTCGACATTCCGATCTCATGCGAGCAGGGGATCTGTGGGACGTGCGTTATGAAAGTCT<br>TAGACGGCGAACCGGACCATCGGGATACTTTTTTGACCGATCAAGAGCGGGCTGCAA<br>ACGACCGGTTCACTCCATGTTGTTCACGGAGCAAAACCGCCCGTCTGGTGATTGATTT<br>TTAA |
| *mpGdmA* | ATGGTAGCCGAGACTCAGATCCCGTATCTGACAAACGCATGGTATGTAGCAGCGCTGT<br>CATCGGAGGTTGGCGCCCAGGCCCTGTTCCACCGTAAGATTCTCGATACATCCGTCTT<br>GATTTACCGCAAGCAGGACGGGGAGGCTGTAGCGTTACACGACCGCTGCCCTCACC<br>GTTTCGCCCCACTCCATTTAGGTAAACGCATCGGTGATGAAGTGGCTTGTCTGTACCAT<br>GCCTTGCAATTTGACTGCTCAGGCCAGTGTACGAAGAATCCGCATGGGAACGGTCAG<br>ATCCCGAAGGCCGCAAAAGTTCGGTCATTTCCACTTGTAGAACGGTATGGTTTTCTGTG<br>GATTTGGATGGGCGAGGACGCCCCGGACGTAGCGCGCTTACCTGATTTCGGGGAGC<br>TTGATAATGGGCCTGAGACTGGGGTTGCCTATACCTACATGCACATGAAGGCCAACTA<br>TGAGCTTATCATCGACAATGTGATGGATCTGAGCCACATTGATCATGTACACGGTGAGA<br>TCATCTCAACGCGGGGGCAACTGTCGCCACTGGTGCCACAGATGCGCGAGGGTGAA<br>CGGGCTATCAGCGCCCGTTGGGAGTGGCAGCAAACTCCAGCCATGCTGATCTTCGCA<br>AACTTTTTGCCTGAGCCTGCCGCTGGTGCACGGCATTTTTTCGACATCACTTGGACTCC<br>ACCGGCAAATATCCAACTCTCAGTCGGTGCTACCCAAGATGGTGGTGCCCTCGACCT<br>TGCGGGCTGCATCGGGCAATATGATCTGCACACTTGCACCCCGGAAACGGCGAACA<br>CGACCCATTATTGGTTTGCAACGCGGCGGAACCACGTTGTAGAGGATGCCGATTACAA<br>CGCCATGAAGATTCAAGCAATGCACGCTGCTTTTGAGAATGAGGATGGGCCAATTATC<br>GAAGCCGTCCATGATGAAATGGGTACAACGGATTTCTTTTCCTTAAATCCGGTCCTCAT<br>GACAAATGACGTTGCTCCAGTGAAAGTACGTCGGCGTCTGCGTCAATTAATTTCGTGA |
| *mpGdmB* | ATGGAGGTGACAGTAGCAGGTCGCGAACCGGCCGCAGAAGGGGTGGCCTCATTTGA<br>GCTGCGTCGTGCTGACGGCGCTCCACTCCCTGCGTTTGATGCCGGCGCACATATCGA<br>CGTAGAGTTGCCAAACGGCATGGTTCGTCAGTATAGCCTGTGCAACGACCCGGCCGA<br>GCGTGGGTATTATCGGATTGGCGTACTGCTGGAAGCCGCTGGTCGCGGTGGTTCCCG<br>CTCCGCACACGAGCAGCTGTTGCGGGGCAGCCGCCTTCGGATTAGCGCACCGCGG<br>AATCATTTTGCCTTGGTCCCGGCCCGCCATACTCTGTTATTAGCTGGGGGTATTGGTGT<br>GACACCGATCCTCTCGATGGCTGAACAGCTGGCTCACGCTGACGCGTCTTTTGAATTA<br>CATTATTGTAGCCGCAGCCCTGCACGGACCGCGTTCGTACCGCGGCTCCAGGCAGC<br>CGCGTTTGCTGCTCGCGTACACTTCCATTTCGACGATGGTGATGCTGCGCAACGCTTT<br>GACGCACGTGCCGTGTTGGCGCAAGCTGTAGACACCACACACCTTTATGTGTGTGGC<br>CCTAAAGGTTTCATGGACCATGTAATCCTGACCGCTCGGGCACTGGGCTGGTCAGAA<br>GGTAACATCCATTACGAGTATTTTGCAGGTGCAGCCGTCGACTCTAGTGCGGACCGC<br>GGTTTTGAGGTCCAAATGGCTGGCAGCTCTCAAGTCGTGCCGGTGGCTCCTGGGCAA<br>TCAGTAGTTCAGGCCTTGGCTGCTCGTGGTATTGATGTGCCGGTATCGTGTGAGCAGG<br>GCGTGTGTGGTACCTGTGTGATGCGTGTGGTCCAAGGGGAGCCAGAACATCGCGACA<br>TGTATTTCAGTGCCGCAGAGCATGCAGCCAATGACCGTTTTACTCCATGTTGCTCGCG<br>CAGTAAATCTGCGCGTCTTGTAATTGACTTCTAA |

| | |
|---|---|
| *psGdmA* | ATGGTAGCCAAGTCCAATCCTACATATCTTCTCAATGCCTGGTATGTCGCAGCGTTGTC<br>AACCGAAGTAGGCCCGGAAGCGTTATTCCATCGCACCATTTTAGATACGTCAGTGCTC<br>ATGTACCGTAAACAAGATGGGACTCCGGTAGCGCTGCACAATCGTTGCCCTCATCGC<br>TTCGCCCCGTTGCACCTCGGGACTCGGCACGGCGATGAGGTTGCCTGCCGCTACCA<br>CGCGCTTCAGTTCGACTGTTCAGGCAAATGTACTAAAAATCCACACGGTAATGGCCAT<br>ATCCCACAAGCAGCTCAAGTACGGCAGTTCCCTCTGGTTGAGCGGTTCGGCTTTCTCT<br>GGATCTGGATGGCAGAGGAACAACCGGACTATGAGCGGCTTCCTGATTTTGGTCCACT<br>GGACGAGGGGCACCCGAATGGGATTGGTTATACGTATATGCACATGCCAGTTAATTAT<br>GAGCTCATTATCGATAATGTGATGGATTTGTCTCACATCGACCATGTGCATGGGGAGAT<br>TATTTCCACACGCGGGAAACTTTCTCCGGTAGTTCCTAAAGTTCAGGAACTCGATCGTG<br>CCATTGTCGCTCGGTGGGAGTGGAAACAAACTCCTGCAATGATGATCTTTGCTAACTTT<br>CTTAGTCAACCACGCGAGGAGGCCGATCACTACTTTAATATTACGTGGACTCCTCCGG<br>CCAATATTCAGTTATCCGTGGGGGCTGTTCAAGGCGAGAAGTCCTTCGACGAGTGCAT<br>CGGCCAGTATGACCTCCATACCACTACGCCAGAGAGCCAGTTCAGTACCCACTACTT<br>TTTCGCTACCCGCCGCAACCATAATGAGGAAGACGGTGAGTACAATGCTATTAAGATC<br>AAGGCCATGCACCAGGCCTTCGAAGACGAGGATGGTCCGATCATTCGCGCAGTACA<br>GGATGAAATGGGGACTAGCGAGTTTTTTGAGCTGGACCCGGTTCTTATGTCTAATGATG<br>TCGCACCAGTGAAGGTTCGCCGGTTATTAAAACGCCTCATTGATGACGAGGGTTCGTG<br>A |
| *psGdmB* | ATGCAGGTCCAAGTAACAGCAAAAACGGCTATTGCAGAGGGCGTTATGTTGTTCGAAT<br>TTCGTGCTTCGCCAGAAGACTTGCTTCCACCATTTACTGCAGGTTCCCATATCGACATT<br>GCACTCCCGAATGGGCTCATCCGTCAGTACTCGCTGTGTAATAGTCCTGATGAGCGCA<br>AGCGCTACGAGATCGCTGTATTACTGGAACCAAGTGGGCGTGGCGGGTCCCGTTCCG<br>CACACATTGATTTGCTTCGTGGCGATACGGTGCGTATTAGCGAACCGCGCAATATGTTT<br>CCGCTTGTTAACGCCCGTCACTCGATCTTAGTCGCAGGTGGTATTGGGGTTACACCGA<br>TCTTGTCTATGGCGGAGCACCTTGCCCGTCGTGGCGCATCCTTCGAGCTCCATTACTG<br>TTCCCGTAGCGTAGCTCGGACGGCGTTTCTCGAGCGGATTCAAAGCAGCGCCTTTGC<br>GAACCGGGTCCGCGTATATATTGACGATGATCACGGTCGTGGTAATTTAGATGTGTCC<br>GCAGTTTTACGTGCTCCTTCCCCTGAAACACACCTTTATGTTTGCGGGCCTAAAGGCTT<br>TATGGACTACGTTATCCAGAGCGCCAATGGGGCGGGTTGGGCAGACCGGTGCGTTCA<br>CAAAGAATATTTTGCTGCCGGGACTATTGATGAGGGCAAAGATCGGCCGTTCGAAGTA<br>GTACTCGCAAGTACGGGCCGGGTGGTACCAGTAACGTCTGAGCAGACAGTTGTTCAG<br>GCGCTGGCTCTCCACGGCATCACAGTACCAGTGTCTTGTGAACAGGGTTTCTGTGGTA<br>CATGCGCGATTCGGGTATTGGACGGGGTCCCTGATCATCGGGATGCGTTCTTCTCAGA<br>CGAAGCTAAAGCTGCAAACGATTGCTTCACCCCGTGTTGTTCTCGTGCCAAGTCCGCG<br>CAATTAGTGGTCGATTACTAA |
| *catA* | ATGACCGTGAAAATTAGCCATACCGCAGATATTCAGGCCTTTTTTAACCGTGTTGCAGG<br>TCTGGATCATGCAGAAGGTAATCCGCGTTTTAAGCAGATTATTCTGCGTGTTCTGCAGG<br>ATACCGCACGTCTGATTGAAGATCTGGAAATTACCGAAGATGAATTTTGGCATGCCGTG<br>GATTATCTGAATCGTTTAGGTGGTCGTAATGAAGCAGGTCTGCTGGCAGCCGGTCTGG<br>GTATTGAACATTTTCTGGATCTGCTGCAGGATGCAAAAGATGCCGAAGCAGGTTTAGG<br>CGGTGGTACACCGCGTACCATTGAAGGTCCGCTGTATGTTGCGGGTGCACCGCTGGC<br>ACAGGGTGAAGCACGTATGGATGATGGCACCGATCCGGGTGTTGTTATGTTTCTGCAG<br>GGTCAAGTTTTTGATGCAGATGGTAAACCTCTGGCAGGCGCAACCGTTGATCTGTGGC<br>ATGCAAATACCCAGGGCACCTATAGCTATTTTGATAGCACCCAGAGCGAATTTAATCTG<br>CGTCGTCGTATTATTACCGATGCGGAAGGTCGTTATCGTGCACGTAGCATTGTTCCGA<br>GCGGTTATGGTTGTGATCCGCAGGGTCCGACACAAGAATGTCTGGACCTGCTGGGTC<br>GTCATGGTCAGCGTCCGGCACATGTTCATTTTTTCATTAGCGCACCGGGTCATCGTCAT |

| | |
|---|---|
| | CTGACCACACAGATTAACTTTGCCGGTGATAAATATCTGTGGGATGATTTTGCCTATGC<br>AACCCGTGATGGTCTGATTGGTGAACTGCGTTTTGTTGAAGATGCAGCAGCAGCACGT<br>GATCGTGGTGTTCAGGGTGAACGTTTTGCAGAACTGAGCTTTGATTTTCGCCTGCAGGG<br>TGCCAAAAGTCCGGATGCAGAAGCCCGTAGCCATCGTCCGCGTGCACTGCAAGAAG<br>GGAATTCATGA |
| *bcER* | ATGGTAGCCAAATACAAGAAACTGTTCGAAACCGTGAAAATCCGTAACGTGGAACTGA<br>AAAATCGTTATGCAATGGCACCGATGGGTCCGCTGGGTTTAGCAGATGCAGAAGGTG<br>GTTTTAATCAGCGTGGTATTGAGTATTATACCGCACGTGCCCGTGGTGGCACCGCACT<br>GATTATTACCGGTGTTACCTTTGTTGATAACGAGGTTGAAGAACATGGTATGCCGAATGT<br>TCCGTGTCCGACACATAATCCGGTTCATTTTGTTCGTACCAGCAAAGAAATGACCGAA<br>CGTATTCATGCATACGATAGCAAAATCTTTCTGCAGATGAGCGCAGGTTTTGGTCGTGT<br>TACCATTCCGACCAATCTGGGTGAATATCCGCCTGTTGCACCGAGTCCGATTCCGCAT<br>CGTTGGCTGGATAAAACCTGTCGTGAACTGACCGTTGAAGAAATTCATAGCATTGTGC<br>GCAAATTTGGTGATGGTGCATTTAATGCAAAACGCGCAGGCTTTGATGGTGTTCAGATT<br>CATGCAGTTCATGAAGGTTATCTGCTGGATCAGTTTGCAATCGCCTTTTTTAACAAACGT<br>ACCGATGCCTATGGTGGACCGCTGGAAAATCGTCTGCGTTTTGCCCGTGAAATTGTGG<br>AAGAAATTAAACAGCGTTGCGGTGAAGATTTTCCGGTTACACTGCGTTTTAGTCCGAAA<br>AGCTTTATCAAAGATTGGCGTGAAGGTGCACTGCCTGGTGAAGAATTTGAAGAAAAAG<br>GTCGTGATCTGGATGAAGGTATTGAAGCAGCAAAACTGCTGGTTAGCTATGGTTATGAT<br>GCACTGGATGTTGATGTTGGCAGCTATGATAGTTGGTGGTGGTCACATCCGCCTATGTA<br>TCAGAAAAAAGGTCTGTATATTCCGTATGCGCGTCTGGTTAAAGAAGCAGTTGACGTTC<br>CGGTTCTGTGTGCAGGTCGTATGGATAATCCGGATCTGGCACTGGCAGCACTGGAAG<br>ATGGTGCCTGTGATATTATCAGCCTGGGTCGTCCGCTGCTGGCCGATCCTGATTATGTT<br>AATAAACTGCGTATTGGTCAGGTGGCAGATATTCGTCCGTGTCTGAGCTGTCATGAAG<br>GCTGTATGGGTCGTATTCAAGAATATTCAAGCCTGGGTTGTGCAGTTAATCCGGCAGCA<br>TGTCGTGAAAAAGAAGCCGCACTGACACCGGCACTGAAAAAAAAGCGTGTTCTGATTG<br>CCGGTGGTGGTGTTGCAGGTTGTGAAGCAGCCCGTGTTCTGGCCCTGCGTGGTCATG<br>AACCGGTGATTTTTGAAAAAAGCAATCGCTTAGGTGGCAATCTGATTCCTGGTGGCGC<br>ACCGGATTTTAAAGAAGATGATCTGGCGCTGGTTGCATGGTATGAACATACCCTGGAA<br>CGTCTGGGTGTTGAAATTCATCTGAATACAGCACTGACCAAAGAAGAAATCCTGGCAG<br>CAAATGTTGATGCAGTGCTGATTGCAACCGGTAGTAATCCGAAAATTCTGCCGCTGGAT<br>GGTAAAAACAAAGTGTTTACCGCAGAAGATGTTCTGCTGGACAAAGTTGATGCCGGTC<br>AGCATGTTGTTATTGTTGGTGGCGGTCTGGTTGGTTGTGAACTGGCCCTGAATCTGGCC<br>GAAAAAGGTAAAGATGTTAGCCTGGTTGAAATGCAGGATAAACTGTTAGCAGTTAATGG<br>TCCGCTGTGTCATGCAAATAGCGATATGCTGGAACGCCTGGTTCCGTTTAAAGGTGTTC<br>AGGTTTATACCTCCAGCAAAATTGTTGATACCACCGAAAAAACCGCAGTTGTTGATGTG<br>GATGGTGAACTGCGCGAAATTGAAGCCGATAGTATTGTTCTGGCAGTTGGTTATAGCG<br>CAGAGAAAAGCCTGTATGAAGATCTGAAATTTGAAGTGGCCGATCTGCATGTTGTGGGT<br>GATGCACGTAAAGTTGCCAATATTATGTATGCAATCTGGGATGCCTATGAAGTGGCAGC<br>CAATCTTTCGTGA |
| *frmA* | ATGAAATCACGTGCTGCCGTTGCATTTGCTCCCGGTAAACCGCTGGAAATCGTTGAAA<br>TTGACGTTGCACCACCGAAAAAAGGTGAAGTGCTAATTAAAGTCACCCATACCGGCGT<br>TTGCCATACCGACGCATTTACCCTCTCCGGCGATGACCCGGAAGGTGTATTCCCGGT<br>GGTTCTCGGTCACGAAGGGGCCGGCGTTGTGGTTGAAGTCGGTGAAGGCGTAACCAG<br>CGTCAAACCTGGCGACCATGTGATCCCGCTTTACACCGCGGAGTGCGGCGAGTGTGA<br>GTTCTGTCGTTCTGGCAAAACTAACCTCTGTGTTGCGGTTCGCGAAACCCAGGGTAAA<br>GGCCTGATGCCAGACGGCACCACCCGTTTTTCTTACAACGGGCAGCCGCTTTATCAC<br>TACATGGGGTGCTCTACATTCAGTGAATACACCGTAGTCGCGGAAGTGTCTCTGGCAA |

| | |
|---|---|
| | AAATTAATCCAGAAGCAAACCATGAACACGTCTGCCTGCTGGGCTGTGGCGTGACCA<br>CCGGTATTGGCGCGGTACACAACACAGCTAAAGTCCAGCCAGGTGATTCTGTTGCCG<br>TGTTTGGTCTTGGCGCGATTGGTCTGGCAGTGGTTCAGGGCGCGCGTCAGGCGAAAG<br>CGGGTCGGATTATCGCTATCGATACCAACCCGAAGAAATTCGATCTGGCGCGTCGCTT<br>CGGTGCTACCGACTGCATTAACCCGAATGACTACGACAAACCGATAAAAGATGTCCTG<br>TTGGATATCAACAAATGGGGTATCGACCATACCTTTGAATGCATCGGTAACGTCAACGT<br>GATGCGTGCGGCGCTGGAAAGTGCGCACCGCGGCTGGGGTCAGTCGGTGATCATCG<br>GGGTCGCGGTTGCCGGTCAGGAAATCTCCACCCGTCCATTCCAGTTGGTCACTGGTC<br>GCGTATGGAAAGGTTCCGCGTTTGGCGGCGTGAAAGGTCGTTCCCAGTTACCGGGCA<br>TGGTTGAAGATGCGATGAAAGGTGATATCGATCTGGAACCGTTTGTCACGCATACCAT<br>GAGCCTGGATGAAATTAATGACGCCTTCGACCTGATGCATGAAGGCAAATCCATTCGA<br>ACCGTAATTCGTTACTGA |
| *frmB* | ATGGAACTCATTGAAAAACATGTCAGCTTTGGCGGCTGGCAAAATATGTATCGGCATTA<br>TTCCCAATCACTGAAATGTGAAATGAATGTCGGCGTCTATCTCCCACCAAAAGCCGCG<br>AATGAAAAATTGCCGGTGCTGTACTGGCTTTCAGGCCTGACCTGCAACGAGCAGAATT<br>TCATTACTAAATCGGGGATGCAGCGTTACGCGGCTGAGCACAACATTATTGTTGTTGCG<br>CCGGACACCAGTCCGCGAGGCAGTCATGTCGCAGATGCTGACCGTTACGATCTCGG<br>GCAAGGTGCCGGGTTTTACCTGAACGCGACGCAAGCGCCGTGGAATGAACATTACAA<br>AATGTATGACTATATCCGCAACGAGCTGCCGGATTTAGTGATGCATCATTTTCCGGCAA<br>CGGCCAAAAAGTCTATCTCTGGTCATTCTATGGGCGGGCTGGGCGCGCTGGTGCTGG<br>CGTTACGTAACCCAGATGAATATGTCAGTGTCTCGGCGTTTTCGCCCATTGTCTCCCCA<br>TCGCAAGTGCCGTGGGGACAGCAAGCCTTTGCTGCATATCTTGCTGAAAATAAAGATG<br>CCTGGTTGGATTACGACCCGGTGAGTCTTATTTCACAAGGTCAACGCGTTGCGGAAAT<br>CATGGTTGATCAGGGGTTGAGTGATGATTTTTACGCAGAACAGCTGCGGACTCCAAAT<br>CTTGAAAAGATCTGCCAGGAGATGAATATCAAGACGTTAATCCGTTATCACGAGGGTTA<br>TGATCACAGCTATTATTTTGTCTCCAGTTTTATTGGCGAGCATATTGCCTACCACGCCAA<br>TAAACTGAATATGCGTTGA |
| *fdhA* | ATGTCAGGGAATAGGGGAGTAGTTTATCTAGGTTCTGGCAAGGTGGAAGTTCAGAAAAT<br>CGATTATCCGAAGATGCAAGATCCGCGTGGTAAAAAGATCGAGCACGGTGTGATCCT<br>GAAAGTTGTCTCGACCAATATCTGCGGCTCTGATCAGCACATGGTTCGTGGTCGTACC<br>ACCGCCCAAGTGGGTTTGGTTCTGGGTCACGAGATCACCGGTGAGGTTATTGAAAAGG<br>GTCGGGACGTGGAGAACCTGCAGATCGGCGAcctgGTTTCCGTGCCGTTTAACGTCGC<br>GTGTGGTCGTTGCCGTAGCTGCAAAGAAATGCACACCGGTGTGTGCTTGACGGTTAAT<br>CCGGCGAGAGCAGGCGGCGCGTACGGTTACGTTGACATGGGTGACTGGACCGGAG<br>GCCAGGCTGAATATCTGCTGGTTCCGTACGCCGACTTTAACCTGTTGAAGCTGCCAGA<br>TCGCGATAAAGCTATGGAAAAGATCCGCGACTTGACGTGCCTGAGCGACATCCTGCC<br>AACGGGTTACCATGGCGCGGTGACTGCGGGTGTTGGTCCGGGTAGCACCGTCTATGT<br>GGCGGGTGCGGGTCCGGTGGGTCTTGCTGCGGCTGCCAGCGCACGCTTGTTAGGTG<br>CCGCGGTTGTGATTGTAGGCGATTTAAACCCGGCCCGCCTGGCCCATGCAAAAGCTC<br>AAGGCTTCGAAATTGCAGACCTCTCCCTGGATACCCCGCTGCACGAGCAGATTGCGG<br>CGCTGTTGGGTGAACCGGAAGTCGACTGTGCAGTTGACGCGGTTGGCTTCGAGGCTC<br>GCGGTCACGGCCACGAGGGTGCGAAGCACGAGGCACCGGCGACCGTGCTCAACAG<br>CCTGATGCAAGTTACCCGTGTGGCCGGCAAGATCGGCATTCCGGGCCTGTATGTTAC<br>GGAAGACCCGGGTGCAGTGGACGCGGCGGCGAAGATCGGCTCTCTGAGTATTCGTTT<br>TGGTCTGGGCTGGGCGAAGTCCCATAGCTTTCATACTGGTCAAACCCCTGTGATGAAA<br>TACAATCGTGCTTTGATGCAGGCGATTATGTGGGATCGTATTAACATCGCGGAGGTAGT<br>TGGCGTCCAGGTGATTAGCCTGGATGATGCTCCGCGTGGCTACGGCGAGTTCGATGC<br>AGGCGTGCCGAAGAAATTCGTCATCGACCCGCATAAAACCTTCAGCGCTGCCTAA |

| | |
|---|---|
| *GroES/EL*<br>JUMP<br>parts | ATGAATATTCGTCCATTGCATGATCGCGTGATCGTCAAGCGTAAAGAAGTTGAAACTAA<br>ATCTGCTGGCGGCATCGTTCTGACCGGCTCTGCAGCGGCTAAATCCACCCGCGGCG<br>AAGTGCTGGCTGTCGGCAATGGCCGTATCCTTGAAAATGGCGAAGTGAAGCCGCTGG<br>ATGTGAAAGTTGGCGACATCGTTATTTTCAACGATGGCTACGGTGTGAAATCTGAGAAG<br>ATCGACAATGAAGAAGTGTTGATCATGTCCGAAAGCGACATTCTGGCAATTGTTGAAGC<br>GTAATCCGCGCACGACACTGAACATACGAATTTAAGGAATAAAGATAATGGCAGCTAA<br>AGACGTAAAATTCGGTAACGACGCTCGTGTGAAAATGCTGCGCGGCGTAAACGTACT<br>GGCAGATGCAGTGAAAGTTACCCTCGGTCCAAAAGGCCGTAACGTAGTTCTGGATAAA<br>TCTTTCGGTGCACCGACCATCACCAAAGATGGTGTTTCCGTTGCTCGTGAAATCGAAC<br>TGGAAGACAAGTTCGAAAATATGGGTGCGCAGATGGTGAAAGAAGTTGCCTCTAAAGC<br>AAACGACGCTGCAGGCGACGGTACCACCACTGCAACCGTACTGGCTCAGGCTATCAT<br>CACTGAAGGTCTGAAAGCTGTTGCTGCGGGCATGAACCCGATGGACCTGAAACGTGG<br>TATCGACAAAGCGGTTACCGCTGCAGTTGAAGAACTGAAAGCGCTGTCCGTACCATG<br>CTCTGACTCTAAAGCGATTGCTCAGGTTGGTACCATCTCCGCTAACTCCGACGAAACC<br>GTAGGTAAACTGATCGCTGAAGCGATGGACAAAGTCGGTAAAGAAGGCGTTATCACC<br>GTTGAAGACGGTACCGGTCTGCAGGACGAACTGGACGTGGTTGAAGGTATGCAGTTC<br>GACCGTGGCTACCTGTCTCCTTACTTCATCAACAAGCCGGAAACTGGCGCAGTAGAA<br>CTGGAAAGCCCGTTCATCCTGCTGGCTGACAAGAAAATCTCCAACATCCGCGAAATG<br>CTGCCGGTTCTGGAAGCTGTTGCCAAAGCAGGCAAACCGCTGCTGATCATCGCTGAA<br>GATGTAGAAGGCGAAGCGCTGGCAACTCTGGTTGTTAACACCATGCGTGGCATCGTG<br>AAAGTCGCTGCGGTTAAAGCACCGGGCTTCGGCGATCGTCGTAAAGCTATGCTGCAG<br>GATATCGCAACCCTGACTGGCGGTACCGTGATCTCTGAAGAGATCGGTATGGAGCTG<br>GAAAAAGCAACCCTGGAAGACCTGGGTCAGGCTAAACGTGTTGTGATCAACAAAGAC<br>ACCACCACTATCATCGATGGCGTGGGTGAAGAAGCTGCAATCCAGGGCCGTGTTGCT<br>CAGATCCGTCAGCAGATTGAAGAAGCAACTTCTGACTACGACCGTGAAAAACTGCAG<br>GAACGCGTAGCGAAACTGGCAGGCGGCGTTGCAGTTATCAAAGTGGGTGCTGCTACC<br>GAAGTTGAAATGAAAGAGAAAAAAGCACGCGTTGAAGATGCCCTGCACGCGACCCGT<br>GCTGCGGTAGAAGAAGGCGTGGTTGCTGGTGGTGGTGTTGCGCTGATCCGCGTAGCG<br>TCTAAACTGGCTGACCTGCGTGGTCAGAACGAAGACCAGAACGTGGGTATCAAAGTT<br>GCACTGCGTGCAATGGAAGCTCCGCTGCGTCAGATCGTATTGAACTGCGGCGAAGAA<br>CCGTCTGTTGTTGCTAACACCGTTAAAGGCGGCGACGGCAACTACGGTTACAACGCA<br>GCAACCGAAGAATACGGCAACATGATCGACATGGGTATCCTGGATCCAACCAAAGTA<br>ACTCGTTCTGCTCTGCAGTACGCAGCTTCTGTGGCTGGCCTGATGATCACCACCGAAT<br>GCATGGTTACCGACCTGCCGAAAAACGATGCAGCTGACTTAGGCGCTGCTGGCGGTA<br>TGGGCGGCATGGGTGGCATGGGCGGCATGA |
